# Discrete and sequential critical periods organise the development of task-specific sensorimotor circuits in mice

**DOI:** 10.1101/2025.09.18.676788

**Authors:** Laura Andreoli, Antonia Maria Constantinescu, Anastasia Garai, Christopher J. Black, Stephanie C. Koch

## Abstract

Somatosensory circuits in early life must maintain stable, task-selective pathways while behavioural repertoires undergo rapid change. How such circuits construct these behaviours under evolving functional demands has remained unclear. Here we show that sensorimotor behaviours are shaped through sequential, experience-dependent critical periods rather than a single global window of plasticity. Using transient perturbations of somatosensory input across postnatal development in mice, we identify three discrete life stages that exert lasting effects on adult behaviour. Perturbation during postnatal days 8-12 selectively increases adult sensitivity to dynamic touch. The same manipulation during days 13-17 produces persistent deficits in motor coordination. Perturbation during days 18-22 instead results in lifelong impairments in skilled locomotion. These findings reveal that somatosensory circuits undergo multiple phases of refinement, each aligned with the changing functional needs of the developing organism. This framework of sequential, task-specific critical periods offers a new model for building lifelong sensorimotor function.

**Significance Statement:** Developing sensory systems must construct precise neural circuits to support dynamic behaviours that change with postnatal experience. We show that somatosensory circuits achieve this through sequential, task-specific critical periods, rather than a single fixed window of plasticity. This dynamic framework demonstrates how experience can guide the stepwise construction of sensorimotor behaviours, offering a new perspective on critical periods across complex, multimodal systems.

## Introduction

Sensory circuits and their evoked responses mature progressively over early postnatal life. During this period, sensory experience plays a pivotal role in shaping the structural and functional organisation of neural networks, establishing the foundations for modality-specific behaviours that persist throughout life (Micheva and Beaulieu 1995; Waldenstrom, Christensson, and Schouenborg 2009; Koch and Fitzgerald 2013; Hubel and Wiesel 1970; Carvell and Simons 1996; Chang and Merzenich 2003; Franks and Isaacson 2005). In many sensory systems, this experience dependent refinement is focused into critical periods, transient windows of heightened plasticity during which sensory input can leave lasting imprints on circuit architecture. These critical periods are well characterised in visual, auditory, and olfactory systems (Franks and Isaacson 2005; Hensch 2004; Hubel and Wiesel 1970), but their timing, structure and functional role in the somatosensory system remain less well defined.

Somatosensation is unique among sensory modalities in that it encompasses multiple distinct submodalities, including touch, nociception, proprioception, and itch, each driving specific reflexive behaviours essential for survival (Koch 2019). However, the architectural precision required for neural circuits to recruit task-selective behaviours is not present at birth. Early sensorimotor reflexes are broad, exaggerated, and poorly tuned, reflecting a mismatch between immature primary afferent input and underdeveloped spinal circuits (Waldenstrom, Christensson, and Schouenborg 2009). Adult-like task-specific behaviours instead emerge gradually over the first three postnatal weeks, in parallel with the refinement of afferent terminations, maturation of local sensorimotor circuits, and the arrival of descending corticospinal inputs (Petersson, Granmo, and Schouenborg 2004; Beggs et al. 2012; Waldenstrom, Christensson, and Schouenborg 2009; Martin 2005). This postnatal refinement of behaviour raises a fundamental question: how can task-specific circuits be built when the tasks themselves are changing? One possible solution is the use of sequential critical periods: multiple, temporally distinct windows of plasticity, each aligned with the emergence of new behavioural demands. In this framework, each phase of circuit refinement consolidates the behaviours most relevant to that stage of development, while preserving the capacity for further adaptation as the organism matures.

Here, we test the hypothesis that somatosensory development is governed by multiple, experience-dependent critical periods. Using transient perturbations of somatosensory input in mice, we identify three discrete postnatal windows: neonatal ages postnatal day (P) 8 to P12, juvenile P13 to P17, and adolescent P18 to P22, each of which produces distinct, lasting effects on adult behaviour. Perturbation during the neonatal period, when Aβ afferents are refined and tactile reflexes emerge (Beggs et al. 2002), leads to increased sensitivity to dynamic touch in adulthood. Disruption during juvenile life, when nociceptive afferents strengthen and coordinated motor behaviours develop (Fitzgerald and Gibson 1984; Feather-Schussler and Ferguson 2016), results in persistent heat responses and motor coordination deficits. Lastly, perturbation in adolescence, during the maturation of corticospinal inputs (Martin 2005), leads to long-term impairments in skilled locomotion.

Together, these findings reveal that somatosensory circuits, unlike other sensory modalities, do not rely on a single fixed window of plasticity. Instead, they undergo sequential, task-specific critical periods, each aligned with the evolving functional requirements of early life. This paradigm provides a new framework for understanding how sensorimotor systems are shaped by experience to support lifelong behavioural flexibility.

## Results

### Early Postnatal Sensorimotor Behaviour and Circuits Are Refined Over Three Developmental Windows

We first asked whether defined periods of sensorimotor circuit maturation could be identified anatomically and behaviourally within the early postnatal period. To do this, we examined the temporal maturation of tactile reflexes, which provide a functional readout of spinal sensorimotor integration. In parallel, we assessed the organisation of low-threshold mechanoreceptive afferents (vGluT1^+^), high-threshold unmyelinated afferents (IB4^+^), and descending corticospinal inputs into the lumbar dorsal horn. Together, these measures identified periods of rapid refinement that may represent heightened developmental sensitivity. Repeated brushing of the hind paw revealed a marked increase in dynamic touch sensitivity between the neonatal (P8-11), juvenile (P14-15) and adolescent (P20) stages, indicating that brush reflexes undergo rapid refinement during this period (Fig. 1a). This behavioural maturation was paralleled by structural reorganisation of primary afferent input to the dorsal horn. At neonatal stages, the superficial dorsal horn displayed dense vGluT1^+^ terminal labelling (Fig. 1b), reflecting an early over-representation of low-threshold mechanoreceptive inputs (Xu et al. 2024). By the third postnatal week (P13-17), this vGluT1^+^ extension had largely retracted, coinciding with the maturation of IB4^+^ nociceptive terminals (Fig. 1c) and C fibre synaptic strengthening within the dorsal horn (Fitzgerald and Gibson 1984). This shift in afferent composition suggests a transition from early spinal tactile bias toward the central integration of nociceptive input.

**Fig. 1.**
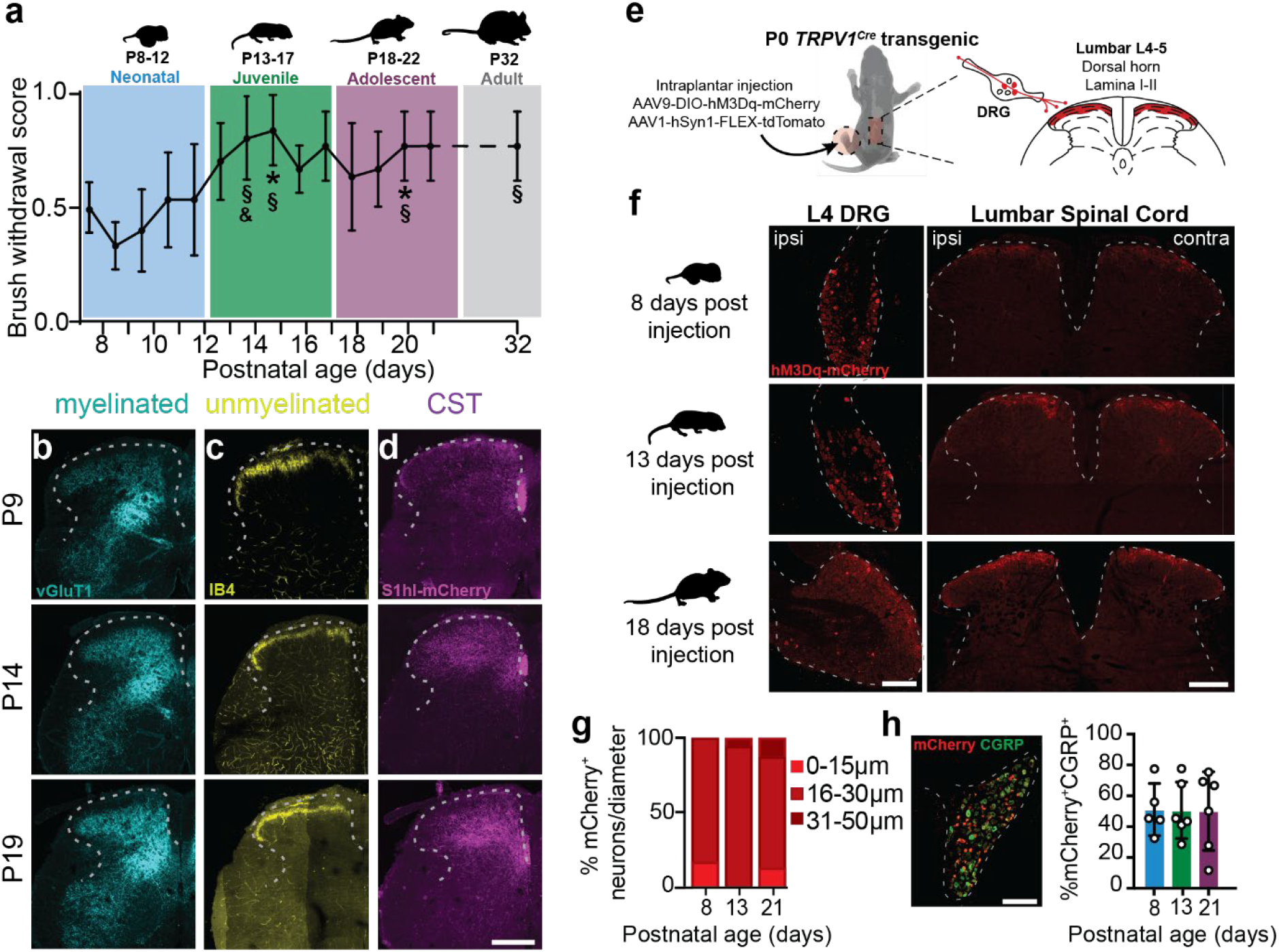
Three sequential windows of sensorimotor plasticity revealed by behaviour and anatomy. (a) Average brush response to five consecutive brush strokes across postnatal development. A significant increase in dynamic tactile sensitivity is observed between the second and third postnatal weeks, indicating that reflexive responses to brush stimulation mature by the end of the second postnatal week (RM one-way ANOVA, *P* < 0.01; Tukey’s test: P8 vs.. P15, P20 \**P* < 0.05; P9 vs.. P14, P15, P20 §*P* < 0.05; P11 vs.. P14 &*P* < 0.05; n = 6). Coloured bars indicate putative developmental windows of plasticity as defined by behaviour and anatomical markers in b-d: blue, neonatal (P8-12); green, juvenile (P13-17); purple, adolescent (P18-22); grey, adult. (b) Representative spinal cord sections immunostained for myelinated afferents (vGluT1^+^, cyan) at P9 (top), P13 (middle), and P18 (bottom). (c) Representative spinal sections immunostained for unmyelinated afferents (IB4^+^, yellow) at the same ages. (d) Representative spinal sections following retrograde mCherry tracing from the somatosensory cortex (magenta), showing progressive targeting of corticospinal input to superficial and deep dorsal horn over postnatal development. These data reveal three developmental windows of central afferent maturation. (e) Schematic of the viral transfection strategy: TRPV1^+^ fibres were selectively targeted by intraplantar injection of a Cre-inducible AAV9-hM3Dq-mCherry vector in P0 *TRPV1*^*Cre*^ pups. (f) Representative images showing hM3Dq-mCherry expression in ipsilateral L4 dorsal root ganglia (DRG) and spinal dorsal horn at (top to bottom) 8, 13, and 18 days post-injection. (g) Quantification of mCherry-labelled DRG soma diameters at each developmental stage (L4-L6; n = 3 per age). (h) Representative section of DRG co-labelled with mCherry (red) and CGRP (green), with quantification of overlap (L4-L6; n = 3 per age). Scale bars: 200 μm (dorsal horn), 200 μm (DRG).

A third phase of refinement was revealed by tracing descending corticospinal projections to the lumbar cord. Corticospinal inputs are known to modulate tactile sensitivity (Liu et al. 2018) and to support the precision of skilled movement in the adult (Martin et al. 2004). In agreement with previous work, we observed a gradual increase in corticospinal terminal density and targeting during the third postnatal week (Martin 2005; Hsu, Stein, and Xu 2006; Donatelle 1977), reaching a mature pattern between P18 and P21 (Fig. 1d). This period coincides with the emergence of skilled motor behaviours (Altman and Sudarshan 1975; Fox 1965; Feather-Schussler and Ferguson 2016) and suggests an additional window of circuit plasticity, distinct from those defined by primary afferent termination patterns.

Together, these findings define the second, third, and fourth postnatal weeks as sequential periods of rapid refinement in spinal somatosensory circuits. These temporally staggered changes identify discrete windows during which sensory experience is likely to exert long-term influence on sensorimotor function.

### Primary Afferents Can be Targeted in a Temporally Specific Manner

We hypothesised that our identified behavioural windows represented critical periods of plasticity, during which sensory drive would set the long-term threshold for circuit activation. To test this, we needed to transiently and selectively manipulate primary afferent input in a spatiotemporally restricted manner across defined developmental stages.

To access a broad range of developing afferent populations, we used a Cre-dependent chemogenetic approach in *TRPV1*^*Cre*^ mice, taking advantage of the widespread expression of TRPV1 in neonatal afferents (Cavanaugh et al. 2011). Intraplantar injection of a Cre-inducible AAV9-hM3Dq-mCherry vector into the left hind paw at P0 (Fig. 1e) resulted in broad labelling of lumbar DRG (L2-L6) and primary afferent terminals in the lumbar dorsal horn by P8 (Fig. 1f); in line with previous findings (Wang et al. 2018). Labelled DRG were predominantly medium in size (P8: 18.2 ± 1.8 μm; P13: 23.0 ± 4.0 μm; Fig. 1g) and largely co-expressed CGRP (P8: 51.1 ± 16.9%; P13: 50.7 ± 18.3%; P21: 50.1 ± 25.5% Fig. 1h), consistent with a broad phenotypic identity. Functional expression of hM3Dq was confirmed by increased c-Fos labelling in afferent termination zones following systemic CNO administration at each developmental stage (Supplementary Fig. S1).

### Adult Dynamic Touch Sensitivity is Established Within the Second Postnatal Week

Having established the ability to manipulate primary afferents in a temporally precise manner, we next asked whether our identified developmental windows represented critical periods for the establishment of long-term sensorimotor function. To meet this definition, altered sensory activity during a given window should leave lasting behavioural consequences (Belford and Killackey 1980; Wiesel and Hubel 1963; Knudsen 1985; Hensch 2004).

Our first window was identified within the neonatal period (P8-12; Fig. 1), when light touch sensitivity decreases as circuits shift from broad A-fibre driven activity toward modality-specific segregation of A- and C-fibre input (Fitzgerald and Jennings 1999). We therefore hypothesised that manipulating primary afferent activity during this period would alter the establishment of adult responses to dynamic touch (Fig. 2a).

**Fig. 2.**
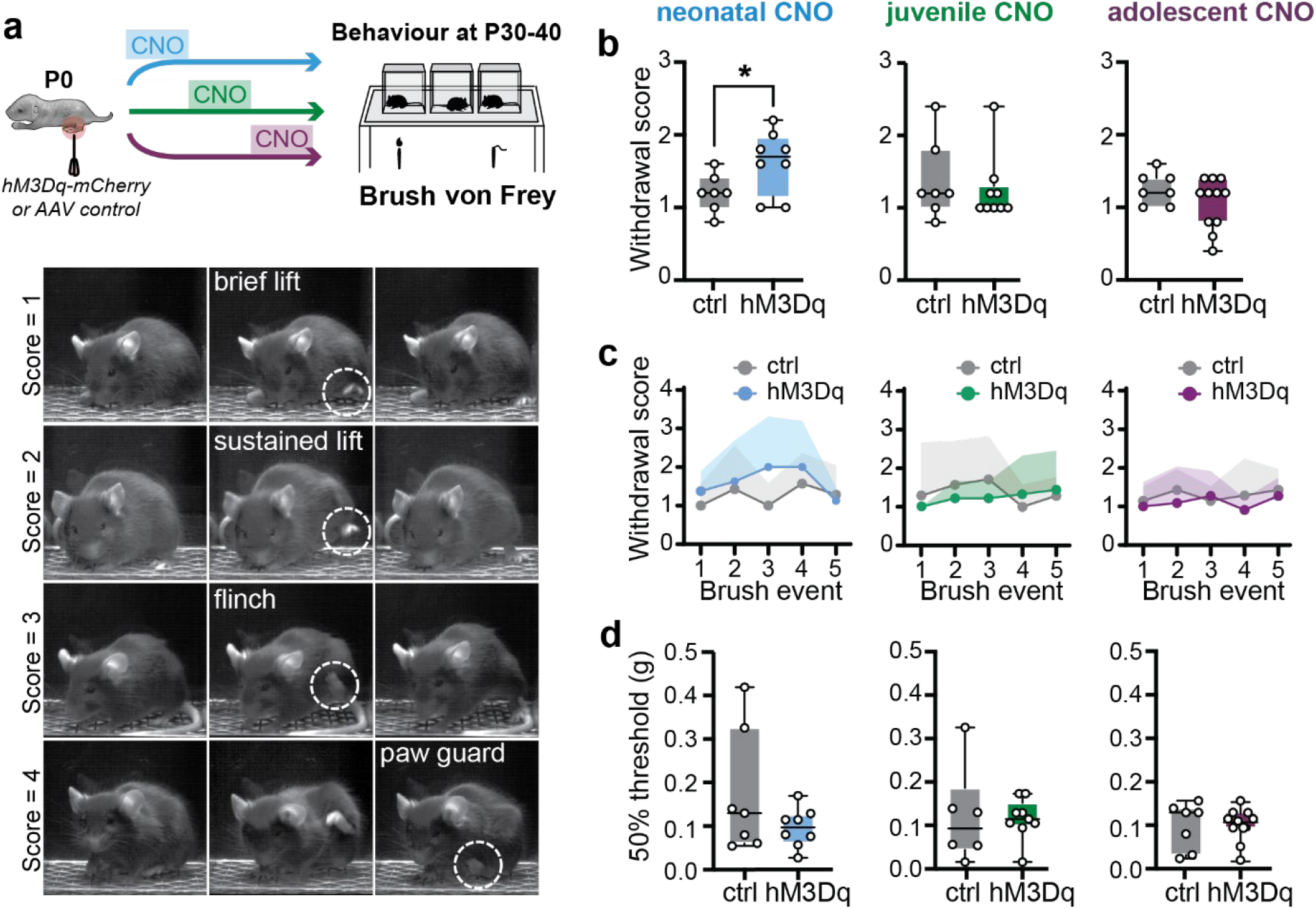
Lasting dynamic touch sensitivity after chronic primary afferent activation in the neonatal period. (a) Schematic (top) of the dynamic brush and static von Frey hair assays, with representative stills (bottom) showing withdrawal responses scored 1-4. (b) Average withdrawal to five consecutive dynamic brush strokes significantly increased in adults with neonatal primary afferent activation (left; P8-12; ctrl vs. hM3Dq, \**P* < 0.05, unpaired t-test), but unchanged in juvenile (middle; P13-17) and adolescent (right; P18-22) groups (both *n*.*s*., Mann-Whitney test and unpaired t-test, respectively). (c) Trial-by-trial analysis revealed no differences across individual brush strokes (all *n*.*s*., RM two-way ANOVA). (d) Static touch sensitivity (von Frey hair thresholds) was unchanged in all groups (all *n*.*s*., unpaired t-tests). Group sizes: neonatal, n = 7 ctrl, 8 hM3Dq; juvenile, n = 6 ctrl, 9 hM3Dq; adolescent, n = 7 ctrl, 11 hM3Dq. *TRPV1*^*Cre*^ mice expressing hM3Dq are shown in blue (neonatal), green (juvenile), and purple (adolescent); controls are shown in grey. All testing was performed in adulthood following transient developmental primary afferent activation.

In adulthood, mice that received neonatal primary afferent activation displayed significantly heightened withdrawal responses to dynamic brush compared to controls (Fig. 2b). This increase was evident when average responses across the five consecutive brush strokes were summed, but not when responses to individual strokes were analysed separately (Fig. 2c), suggesting a long-term increase in dynamic touch behavioural responses rather than altered temporal summation.

In contrast, afferent activation during juvenile (P13-17) or adolescent (P18-22) stages produced no significant changes in adult dynamic brush sensitivity (Fig. 2b-c). Static tactile thresholds measured by von Frey hair were unchanged across all groups (Fig. 2d). No changes in dynamic touch or static tactile thresholds were observed in any group when stimulating the contralateral hind paw (Supplementary Fig. S2).

Together, these results identify the neonatal period (P8-12) as a critical window for establishing dynamic touch sensitivity in adulthood. Perturbation of afferent activity during this period produces a lasting increase in touch reactivity, while later manipulations during juvenile or adolescent stages have no long-term effect.

### Adult *Heat Sensitivity is Established Within the Third Postnatal Week*

Refinement of tactile reflexes secures appropriate responses to innocuous stimuli, but developing circuits must also establish reliable detection of noxious input for survival (Fitzgerald 2005). We therefore examined whether nociceptive responses are shaped within defined developmental windows. Thinly myelinated Aδ-fibres, which mediate responses to sharp mechanical stimuli, project into the dorsal horn around birth and reach their mature termination fields within the first postnatal week (Fitzgerald 2005). In contrast, TRPV1^+^ C-fibres, which transmit nociceptive heat information (Cavanaugh et al. 2009), are present at birth but undergo substantial spinal synaptic strengthening during the second and third postnatal weeks (Fitzgerald and Gibson 1984; Fitzgerald and Jennings 1999). Based on this trajectory, we hypothesised that altering afferent activity during P13-17 would disrupt the consolidation of heat-evoked nocifensive behaviours in adulthood. To test this, we measured both mechanical and thermal nociceptive responses in adult mice following afferent activation during the neonatal (P8-12), juvenile (P13-17), or adolescent (P18-22) windows.

In a ramping hotplate assay (42-50 °C) with an escape platform (Fig. 3a), latency to escape was equivalent across all groups (Fig. 3b), indicating normal detection of thermal stimuli. However, when nocifensive behaviours such as licking, biting, and paw guarding were quantified (Fig. 3c), adult mice that had received juvenile afferent activation displayed a significant reduction in aversive behaviours compared with controls in contrast to neonatal or adolescent groups (Fig. 3d).

**Fig. 3.**
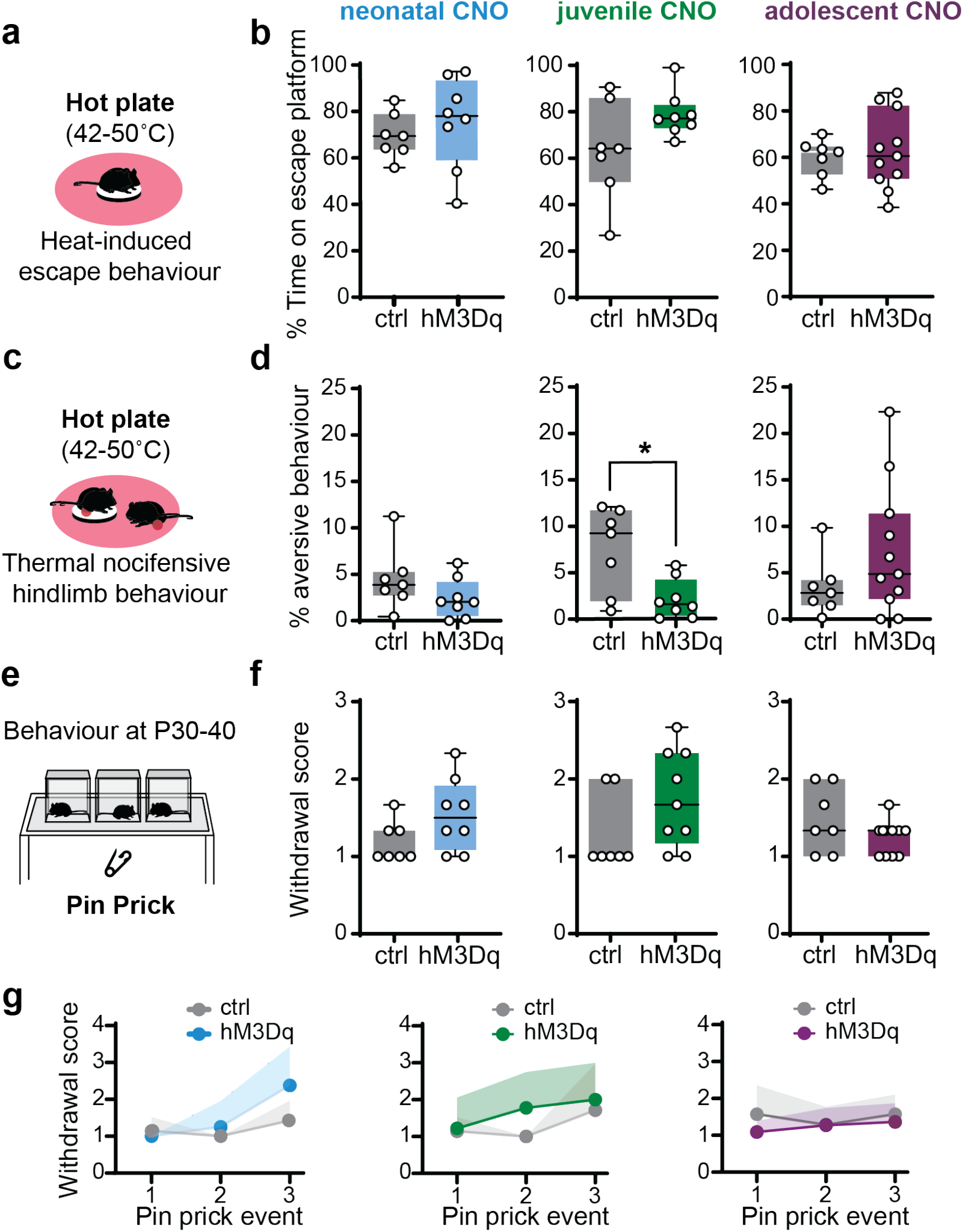
Lasting heat-evoked nocifensive behaviour after chronic primary afferent activation the juvenile period. (a) Schematic of heat-induced escape behavioural assay. (b) Quantification of average heat-induced escape behaviour. Adult mice that had received neonatal (left; P8-12), juvenile (middle; P13-17), or adolescent (right; P18-22) afferent activation showed no differences in escape behaviour (all *n*.*s*., unpaired t-tests). (c) Schematic of thermal nocifensive behavioural assay. (d) Quantification of average heat-induced aversive behaviour. Adults with juvenile afferent activation displayed reduced heat aversive behaviours (middle; ctrl vs. hM3Dq, \**P* < 0.05, unpaired t-test), with no significant effect in neonatal (left) or adolescent (right) groups (all *n*.*s*., Mann-Whitney tests. (e) Schematic of Pin Prick behavioural assay. (f) Quantification of pin prick response. Average withdrawal magnitude to noxious pin prick was unchanged across groups (all *n*.*s*., unpaired t-test or Mann-Whitney test). (g) Response to repeated pin pricks was equally unchanged (all *n*.*s*., RM two-way ANOVA), although neonatally treated mice failed to adapt to repeated stimuli (left; Pin Prick event: F_1.638, 21.30_ = 10.37, *P* < 0.01). Group sizes: neonatal, n = 7 ctrl, 8 hM3Dq; juvenile, n = 6 ctrl, 9 hM3Dq; adolescent, n = 7 ctrl, 11 hM3Dq. *TRPV1*^*Cre*^ mice expressing hM3Dq are shown in blue (neonatal), green (juvenile), and purple (adolescent); controls are shown in grey. All testing was conducted in adulthood following transient developmental primary afferent activation.

Mechanical nociception was assessed using the Pin Prick assay (Fig. 3e). Contrary to that seen in heat behaviours, no significant differences were observed in average pin prick withdrawal magnitude across groups (Fig. 3f). Responses to repeated pin pricks were also unaffected (Fig. 3g), though neonatal animals showed a failure to adapt across trials. No changes in average pin prick withdrawal magnitude or responses to repeated pin pricks were observed in any group when stimulating the contralateral hind paw (Supplementary Fig. S3). Similarly, we observed no changes in cold sensitivity or responses in any group (Supplementary Fig. S4). These findings suggest that early-life afferent activation does not alter the development of mechanical nociceptive reflexes, which may be resistant to transient perturbations in early life.

Together, these findings identify the juvenile period (P13-17) as a critical window for the refinement of heat-evoked nocifensive behaviours. Altered afferent input during this stage selectively weakens the integration of nociceptive signals into appropriate defensive responses, while mechanical nociceptive reflexes remain resilient to perturbation.

### Adult Gross Motor Coordination is Established Within the Third Postnatal Week

The refinement of tactile and nociceptive reflexes ensures appropriate somatosensory responses, but effective interaction with the environment also depends on the emergence of coordinated locomotor patterns. Hindlimb weight support appears near the end of the second postnatal week, and by the beginning of the third week quadrupedal locomotion is reliably expressed, coinciding with refinement of proprioceptive inputs to spinal motor circuits (Feather-Schussler and Ferguson 2016; Martin 2005). We therefore hypothesised that the juvenile period (P13-17), a stage of rapid motor maturation, represents a critical window for consolidating gross locomotor control. To test this, we examined adult locomotor behaviour following afferent activation during the neonatal (P8-12), juvenile (P13-17), or adolescent (P18-22) windows, using the CatWalk XT system to quantify step cycle dynamics (Fig. 4a-b). Neonatal and adolescent groups showed no long-term changes in stance duration (Fig. 4c-d), swing phase duration (Fig. 4e-f), or initial stance (Fig. 4g-h), indicating preserved locomotor development after early or late perturbation. In contrast, juvenile activation produced persistent deficits in adult gait. Mice displayed increased stance duration in the ipsilateral hindlimb (Fig. 4d), reduced swing phase duration (Fig. 4f), and increased initial stance, a measure of postural stability (Fig. 4h), suggesting a deficit in coordination.

**Fig. 4.**
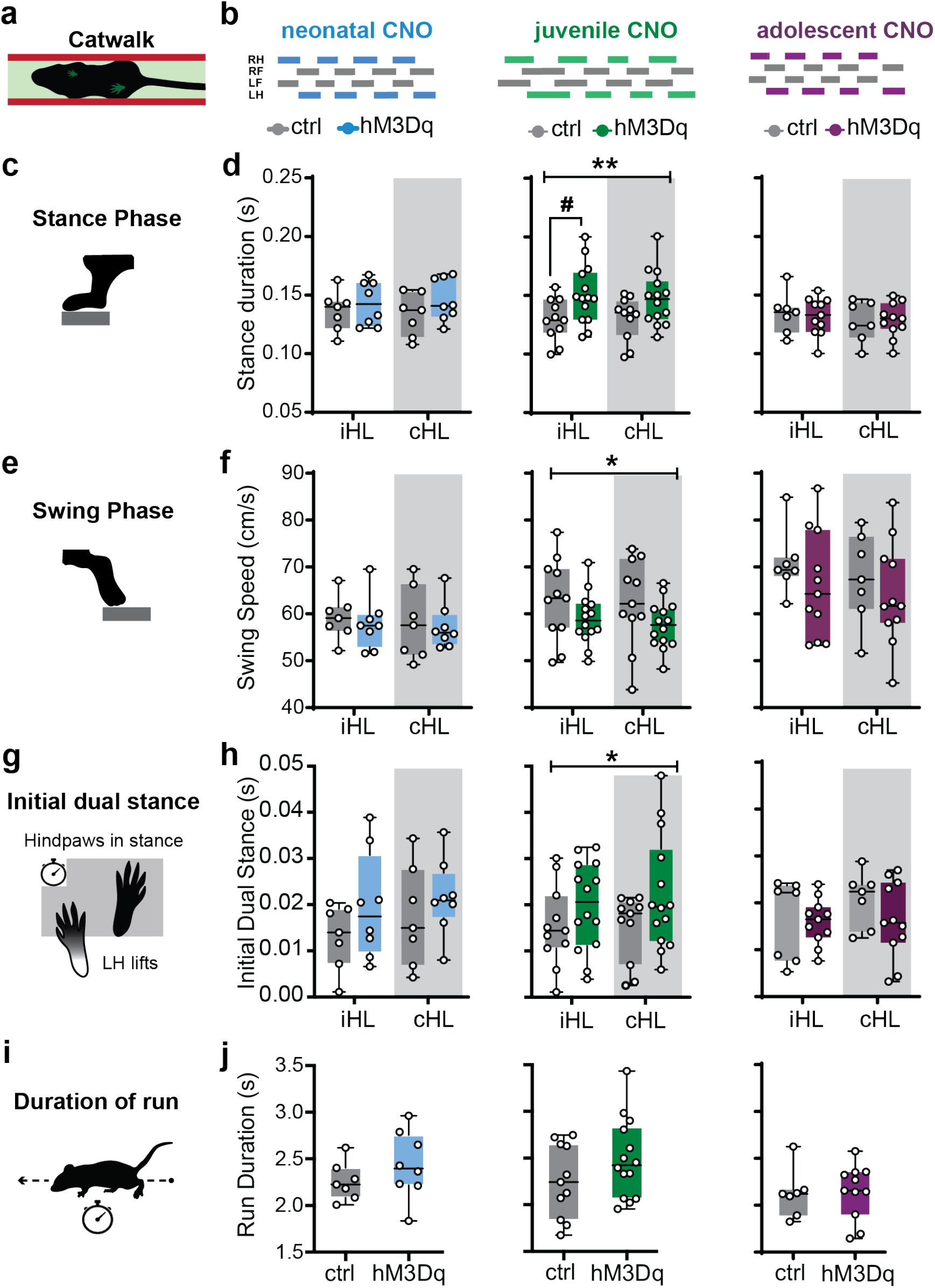
Lasting gross motor coordination deficits after chronic primary afferent activation in the juvenile period. (a) Schematic of CatWalk behavioural assay. (a-b) Representative CatWalk step pattern sequences from adults following neonatal (left; P8-12), juvenile (middle; P13-17), or adolescent (right; P18-22) afferent activation. Solid bars represent stance phase. RH: right hindlimb; RF: right forelimb; LF: left forelimb; LH: left hindlimb. (c) Schematic of hind paw in stance phase. (d) Quantification of stance phase duration across three runs. Average stance phase duration was significantly increased in the ipsilateral hind paw of the juvenile group (middle; F_1,46_ = 9.468, *P* < 0.05; Šídák’s test \**P* < 0.05), but unchanged in neonatal (left) or adolescent (right) groups (both *n*.*s*., two-way ANOVA). (e) Schematic of hind paw in swing phase. (f) Quantification of swing phase duration across three runs. Swing phase duration was similarly altered only in the juvenile group (middle; F_1,46_ = 4.741, *P* < 0.05), with no changes in neonatal (left) or adolescent (right) groups (both *n*.*s*., two-way ANOVA). (g) Schematic of initial dual stance duration. (h) Quantification of initial dual stance duration across three runs. Initial stance was selectively increased in the juvenile group (middle; F_1,46_ = 4.258, *P* < 0.05) but unaffected in neonatal (left) or adolescent (right) groups (both *n*.*s*., two-way ANOVA). (i) Schematic of run duration. (j) Quantification of run duration across three runs. Average run duration was unchanged across all groups (all *n*.*s*. unpaired t-tests). Group sizes: neonatal, n= 7 ctrl, 8 hM3Dq; juvenile, n = 6 ctrl, 9 hM3Dq; adolescent, n = 7 ctrl, 11 hM3Dq. *TRPV1*^*Cre*^ mice expressing hM3Dq are shown in blue (neonatal), green (juvenile), and purple (adolescent); controls are shown in grey. All testing was conducted in adulthood following transient developmental primary afferent activation. Grey shaded areas indicate values contralateral hindlimb values.

Despite changes in coordination, overall locomotor output remained intact: run duration and number of steps were unaffected across all groups (Fig. 4i-j, Supplementary Fig. S5). These findings suggest that juvenile afferent activation does not necessarily impair the ability to generate locomotion, but instead specifically disrupts the fine balance of stance and swing that underpins smooth, coordinated gait.

Together, these results identify the juvenile period as a sensitive window for gross motor coordination, during which peripheral activity shapes the refinement of locomotor circuits.

### Adult Fine Motor Control is Established Within the Third Postnatal Week

Gross locomotion can be generated largely by spinal circuits and does not place strong demands on supraspinal control (Goulding et al. 2014; Grillner 2006; Barbeau and Rossignol 1987). In contrast, skilled locomotion requires integration across the neuraxis, including corticospinal input (Henry et al. 2020; DiGiovanna et al. 2016; Ueno et al. 2018), which matures during the fourth postnatal week (Martin 2005). Having determined that gross locomotor coordination is refined by the third postnatal week, we next asked whether fine motor control is similarly shaped by early sensory experience. To address this, we tested adult mice in the horizontal beam task (Fig. 5a), a voluntary locomotor assay that places high demands on hindlimb precision and corrective reflexes. This paradigm provides a sensitive readout of both fine-scale kinematics and whole-animal performance (Fig. 5b).

**Fig. 5.**
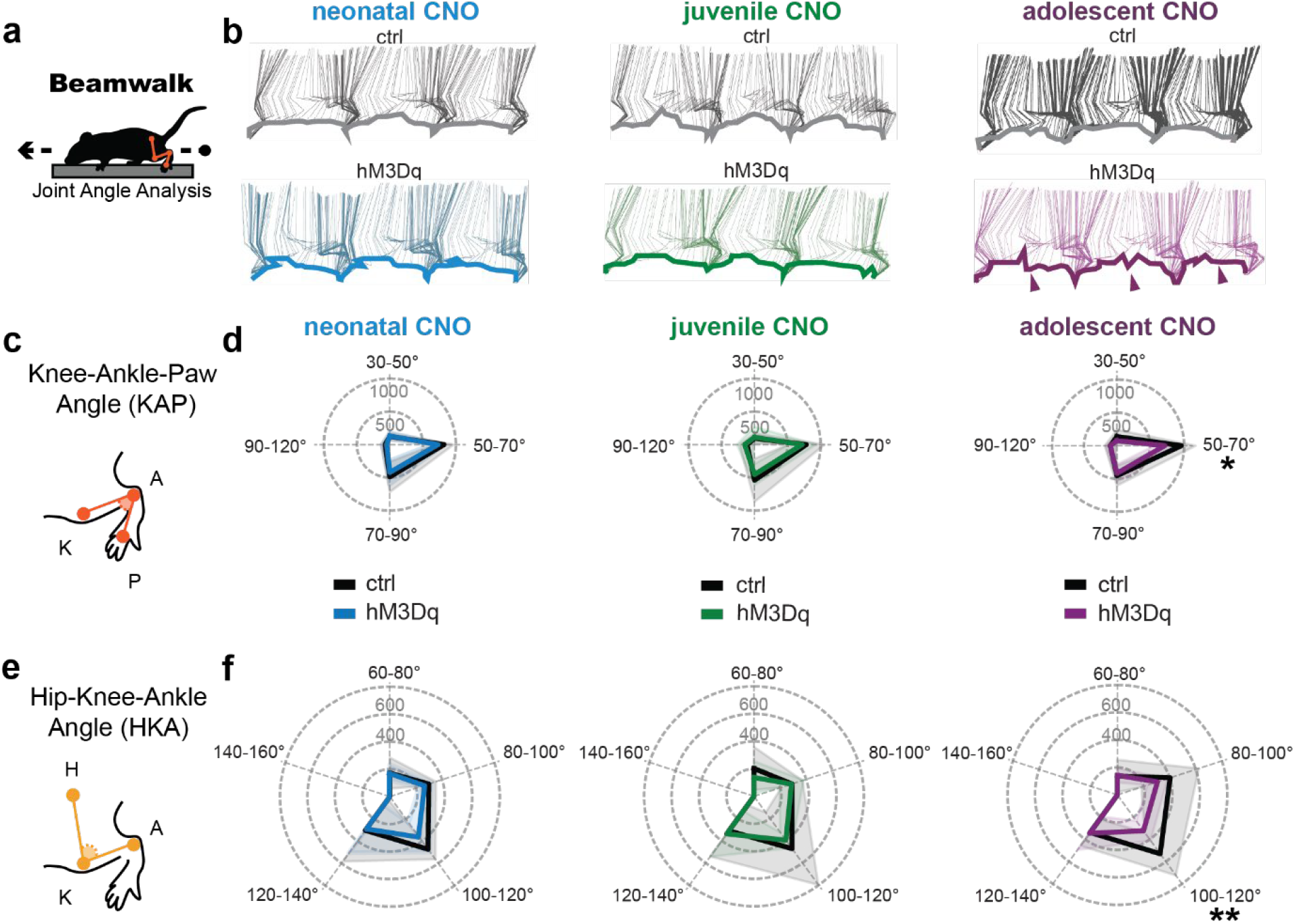
Lasting skilled locomotor impairments after adolescent afferent activation. (a) Schematic of beamwalk behavioural assay. (b) Representative stick figure diagrams showing three complete hindlimb step cycles (swing and stance) for adults following neonatal (left; P8-12), juvenile (middle; P13-17), or adolescent (right; P18-22) afferent activation. Arrows point to loss of swing phase fluidity during stepping. (c) Schematic of knee-ankle-paw (KAP) angle. (d) Spider plot of KAP angle over three consecutive steps. KAP angle was altered only in the adolescent group (right; F_1,60_ = 7.790, *P* = 0.01; interaction F_3,60_ = 3.210, *P* = 0.05), with reductions concentrated between 50-70° (Šídák’s test, \**P* < 0.05). (e) Schematic of hip-knee-ankle (HKA) angle. (f) Spider plot of HKA angle over three consecutive steps. HKA angle was significantly reduced in the adolescent group (right; F_1,75_ = 5.054, *P* < 0.05; interaction F_4,75_ = 2.613, *P* < 0.05), with post hoc analysis showing a selective decrease in the 100-120° range (Šídák’s test, \*\**P* < 0.01). No significant changes were observed in neonatal (left) or juvenile (middle) groups (all *n*.*s*., two-way ANOVA). Group sizes: neonatal, n = 7 ctrl, 8 hM3Dq; juvenile, n = 6 ctrl, 9 hM3Dq; adolescent, n = 7 ctrl, 11 hM3Dq. *TRPV1*^*Cre*^ mice expressing hM3Dq are shown in blue (neonatal), green (juvenile), and purple (adolescent); controls are shown in grey. All testing was conducted in adulthood following transient developmental activation.

Neonatal (P8-12) and juvenile (P13-17) afferent activation did not alter adult skilled locomotion. Hindlimb kinematics were preserved (Fig. 5c-f), and behavioural performance measures such as slips, distance travelled, speed, and run duration were unchanged compared to controls (Supplementary Fig. S6).

In contrast, adolescent (P18-22) activation produced persistent deficits. Adult mice in this group showed reduced knee and ankle joint angles (KAP and HKA) during beam traversal (Fig. 5c-f), with decreases focused within the ranges normally engaged during the swing-stance transition. These kinematic restrictions limited the ability to extend and position the paw securely on the beam. Behaviourally, this was reflected in impaired efficiency: although slips and distance travelled were unaffected, but both speed and run duration were disrupted, indicating less fluent beam crossing (Supplementary Fig. S6).

Together, these findings reveal adolescence as a distinct critical window for skilled locomotion. Peripheral activity during this stage gates not only spinal refinement but also the maturation of cortically gated fine motor control.

### Early-Life Afferent Activation Produces Selective Changes in Dorsal Horn Inhibition

The somatosensory touch and nociceptive phenotypes we observed suggested that transient afferent activation during development might leave lasting imprints on dorsal horn circuits. To test this, we first recorded extracellular activity from the ipsilateral lumbar spinal cord using Neuropixels probes during repeated brush stimulation of the hind paw (Fig. 6a-b). We focused on the neonatal (P8-12) and juvenile (P13-17) groups, which displayed the clearest tactile and gross locomotor phenotypes. We found stimulus-evoked firing of multi-unit activity (MUA) to be preserved in both neonatal and juvenile cohorts: mean z-scored firing rate in the 0-1 s post-stimulus window was comparable between hM3Dq and control animals across all recorded depths (0-500 μm in 100 μm bins; Fig. 6c-d) indicating that sensory responsiveness remains intact after early life altered input.

**Fig. 6.**
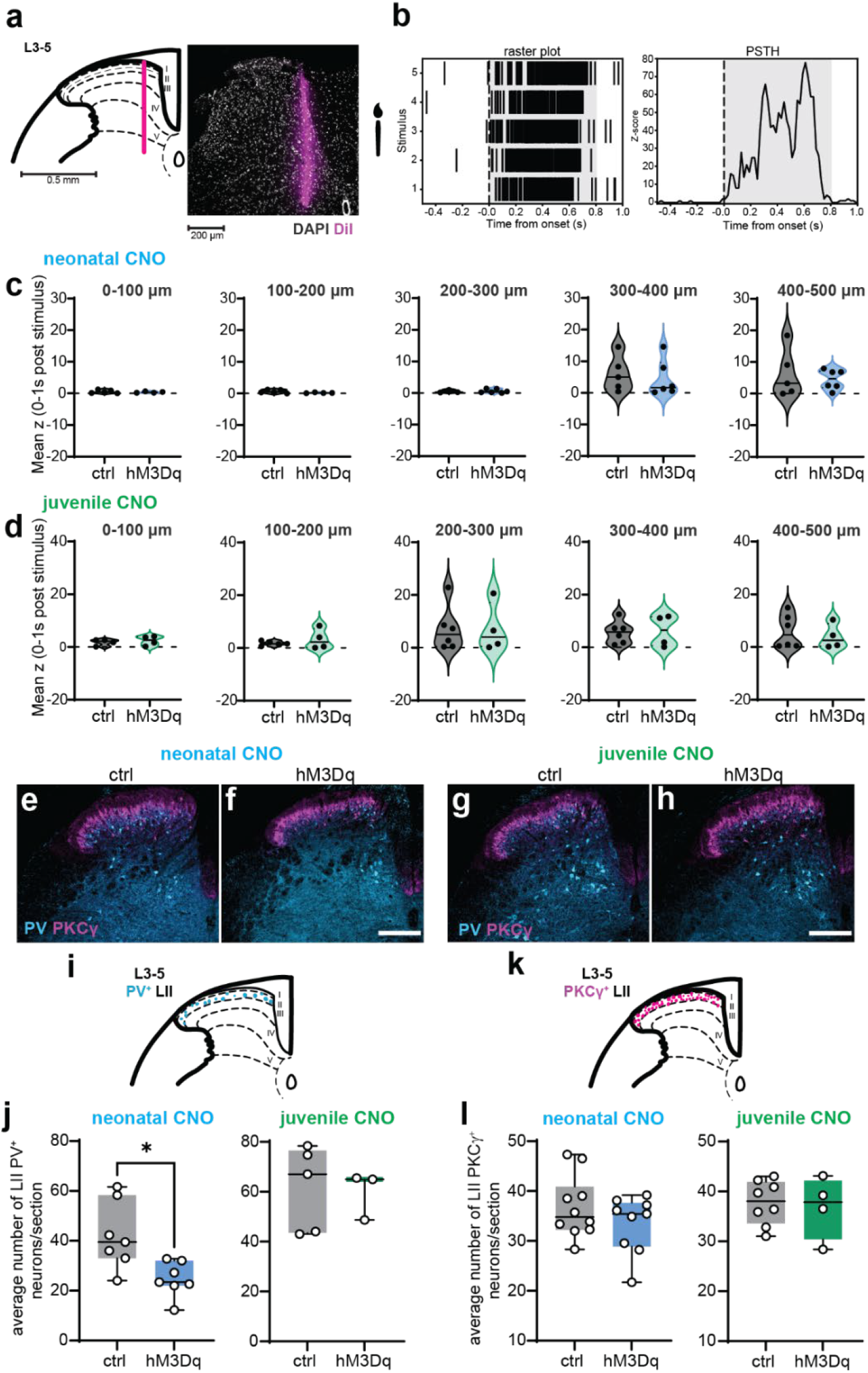
Adult dorsal horn neuronal responses and interneuron populations following early-life afferent manipulation. (a) Schematic of Neuropixels probe insertion into the medial dorsal horn, with confocal image of the ipsilateral hemisection showing DiI stain from the probe track and DAPI counterstain for grey matter visualization. (b) Example spinal MUA response to five consecutive brush stimulations of the ipsilateral hind paw, shown as raster plot (left) and peri-stimulus time histogram (right; PSTH). (c-d) Violin plots showing mean z-scores (0-1 s post-stimulus window) of MUA recorded across probe depth bins (0-100 μm, 100-200 μm, 200-300 μm, 300-400 μm, 400-500 μm) in neonatal (c; P8-12) and juvenile (d; P13-17) groups. No significant differences were observed between control and hM3Dq mice in either group (all *n*.*s*., Mann-Whitney tests). (e-h) Representative images of PV (cyan) and PKCγ (magenta) expression in the ipsilateral dorsal horn of neonatal and juvenile groups. (i) Schematic of localisation of spinal lamina II PV^+^ interneurons used for quantification. (j) Quantification of PV^+^ interneurons in lamina II. PV^+^ interneuron counts were significantly reduced in lamina II of neonatally stimulated mice (left; *P* < 0.05, unpaired t-test) but not in juvenile groups (right; *n*.*s*., Mann-Whitney tests). (k) Schematic of localisation of spinal lamina II PKCγ^+^ interneurons used for quantification. (l) Quantification of PKCγ^+^ interneurons in lamina II. PKCγ^+^ interneuronal counts were unchanged by early life afferent input (all *n*.*s*., Mann-Whitney tests). Group sizes: neonatal, n = 7-10 ctrl, 7-9 hM3Dq; juvenile, n = 5-8 ctrl, 3-4 hM3Dq. *TRPV1*^*Cre*^ mice expressing hM3Dq are shown in blue (neonatal) and green (juvenile); controls are shown in grey. All extracellular recordings and immunohistochemical analyses were conducted in adulthood following transient developmental primary afferent activation. Scale bar: 200 μm.

Whereas MUA firing rate across spinal laminae would inform of the overall output across neuronal phenotypes, we could not gain insight into any alteration in neuronal architecture. We therefore used immunohistochemistry to probe broader circuit organisation (Fig. 6e-h), focusing on interneuron populations known to shape tactile processing. Parvalbumin (PV^+^) interneurons regulate low-threshold tactile gain in lamina II (Petitjean et al. 2015; Boyle et al. 2019), while PKCγ^+^ interneurons relay excitatory touch input from superficial to deeper laminae (Miraucourt, Dallel, and Voisin 2007; Neumann et al. 2008). This pairing allowed us to test whether altered behaviour was associated with any overt changes in inhibitory versus excitatory circuitry. Our analysis revealed a selective reduction of PV^+^ interneurons in lamina II (Fig. 6i-j), with no detectable changes in PKCγ^+^ cell count (Fig. 6k-l). Juvenile animals showed no differences in either population (Fig. 6e-l). No significant changes in primary afferent terminations were observed at either age (Supplementary Fig. S7).

Together, these findings suggest that the consequences of early-life afferent activation are not explained by broad changes in dorsal horn excitability but could instead be the result of selective alterations in inhibitory control. A reduction of PV^+^ interneurons in the neonatal group suggests weakened local gating of tactile input, without corresponding changes in PKCγ^+^ excitatory circuits. This imbalance may amplify brush sensitivity and alter how sensory signals are relayed to deeper laminae for integration with motor output. These results therefore support the view that developmental perturbations do not simply alter somatosensory input but reshape how sensorimotor circuits use sensory drive to guide behaviour.

## Discussion

Our findings reveal somatosensory development to be organised into sequential, task-specific critical periods, during which peripheral activity sets long-term behavioural trajectories. Rather than a single global window of plasticity, we identified three discrete stages: neonatal perturbation increased dynamic touch sensitivity, juvenile perturbation disrupted heat responses and motor coordination, and adolescent perturbation impaired skilled locomotion. Together these results support a framework in which early life experience consolidates behavioural repertoires as they emerge.

This sequential organisation may reflect an adaptive logic: at each stage only the behaviours most relevant for survival are consolidated, preserving flexibility until later functions become essential. Rapid withdrawal reflexes mature during the first three postnatal weeks, by which time they become adult-like in precision and timing (Holmberg and Schouenborg 1996), whereas corticospinal circuits continue to refine into late postnatal development, with functional maturation extending into weeks 5-7 (Martin 2005; Terashima 1995). The narrow time windows we identify (5-6 days each) suggest that somatosensory plasticity is punctuated, with brief bursts of heightened sensitivity interspersed by periods of consolidation, similar to other sensory systems (Hensch 2004).

Across all tested behaviours, transient developmental afferent activation was sufficient to produce long-lasting consequences, demonstrating that peripheral input provides the primary instructive signal for circuit refinement (Fitzgerald 2005). Although TRPV1^+^ afferents are classically associated with nociception (Cavanaugh et al. 2009), their activation during development influenced tactile, nociceptive, and motor outcomes without overt effects on spontaneous behaviours (Supplementary Video V1), suggesting our behavioural effects are unlikely to be the result of stress or injury, but rather a change in spinal afferent drive in early life. This cross-modal impact suggests that it is activity, rather than afferent modality, that determines task-selective circuit maturation.

This experience-dependent maturation appears specific to behaviours requiring inhibitory refinement (touch sensitivity, locomotor coordination) or supraspinal modulation (skilled movement), as opposed to hard-wired protective behaviours such as the nociceptive withdrawal reflex to pin prick or cold (Fig. 3, Supplementary Fig. S4). In agreement with this, our circuit level analyses suggest these alterations in task-selective behaviours are due to changes in inhibitory balance, rather than changes in excitability of dorsal circuits. Spinal extracellular recordings showed preserved stimulus-evoked activity in adulthood, indicating that dorsal horn responsiveness to tactile input remains intact following altered early life experience. However, immunohistochemistry revealed a selective reduction in inhibitory PV^+^ interneurons in lamina II following neonatal activation of afferents, with no change in PKCγ^+^ excitatory neurons. This pattern suggests weakened inhibit rather than altered excitatory relay, consistent with the proposed role of PV^+^ interneurons in tactile gating (Petitjean et al. 2015; Boyle et al. 2019). Such a loss of inhibitory control could lead to increased brush responsiveness, as observed in the neonatally activated animals.

Importantly, our extracellular recordings were targeted to the medial dorsal horn, corresponding to afferent input arising from the hind paw. This maximised the probability of sampling neurons responsive to brushing of the paw but leaves open the possibility that early-life altered experience altered somatotopy, broadening receptive field representations beyond their normal map (Beggs et al. 2002; Granmo, Petersson, and Schouenborg 2008). Our data argue against this explanation. First, we found no significant changes in primary afferent termination patterns in our experimental groups (Supplementary Fig. S7). Second, somatosensory behavioural tests revealed no contralateral effects (Supplementary Fig. S2-S4) and Neuropixels recordings revealed no overt changes in receptive field size between groups (data not shown), suggesting animals retained accurate localisation of paw input. Lastly, our results suggest that early life sensory experience shapes behavioural response magnitude, but not behavioural thresholds, which have been shown to change following receptive field expansion (Beggs et al. 2002). This suggests instead that developmental activity affects how strongly sensory signals are integrated and expressed, rather than whether they are detected at all.

A likely explanation for the sequential emergence of critical periods is that each behavioural repertoire recruits distinct interneuronal ensembles as they come online. Inhibitory interneurons are among the last spinal elements to mature (Baccei and Fitzgerald 2004; Keller et al. 2001; Koch et al. 2012), leaving networks sensitive to altered activity. Distinct populations likely underlie different phenotypes: PV^+^ interneurons regulate tactile gain, glycinergic and dI6-derived interneurons contribute to locomotor patterning (Koch et al. 2017), and corticospinal-responsive populations support skilled movement (Martin et al. 2004). Perturbation during a given window may bias these ensembles into an immature configuration that becomes “locked in.” Recent studies have identified the key role of astrocytes in maintaining inhibitory function and their involvement within critical periods across sensory systems notably following early life injury (Yoo et al. 2025). One hypothesis may therefore be that early life glial function facilitates the maturation of spinal inhibition to regulate the timing and outcome of these somatosensory critical periods. Although outside of the remit of our current study, this remains a key point of interest for future studies.

These findings expand the classical view of critical periods. Whereas vision and audition are defined by a single, well-demarcated window of plasticity (Hubel and Wiesel 1970; Knudsen 1985), somatosensory development proceeds through multiple sequential windows, each tied to the emergence of new functions. This distributed organisation suggests that plasticity is partitioned in time and function, rather than concentrated in a single event. Clinically, this framework implies that early-life injury, inflammation, or abnormal sensory drive, such as neonatal surgery or neuropathy, could bias circuits towards lifelong sensory or motor dysfunction (Walker et al. 2009; Walker 2014; Eccleston et al. 2008; Eccleston and Clinch 2007; Fitzgerald 2024). Our results also suggests that once a developmental window closes, circuits become relatively resistant to later plasticity, helping explain the limited efficacy of adult rehabilitation after injury compared to the flexibility available in early life (Eccleston and Clinch 2007). The therapeutic implication is that the developmental windows we identify may represent opportunities to intervene in children undergoing surgery or other early-life insults to prevent long-term dysfunction.

Together, these findings support a model in which peripheral activity sculpts sensorimotor behaviours across multiple critical periods, each aligned with the evolving demands of development. This sequential architecture allows flexibility while ensuring that essential functions are consolidated in turn, providing a foundation for lifelong behavioural repertoires.

## Methods

### Animals

All experiments were performed in compliance with the United Kingdom Animals (Scientific Procedures) Act of 1986. Animals had ad libitum access to water and food and were housed under 12 hour (hr) light/dark cycles.

Male and female C57BL6 mice (Charles River) were used for the longitudinal dynamic brush study (n=6). *Emx1*^*Cre*^ mice (B6.129S2-Emx1tm1(cre)Krj/J, stock #005628, Jackson Laboratory (Gorski et al. 2002)), were used to trace the developing corticospinal projections from somatosensory cortex hindlimb region (n=3). Male and female *TRPV1*^*Cre*^ mice (B6.129-Trpv1tm1(cre)Bbm/J, stock #017769, Jackson Laboratory) were used for all remaining experiments. Mouse pups in the age groups P8-12 (neonatal), P13-17 (juvenile), and P18-22 (adolescent) were housed with their mothers and littermates until weaned at P22 when they were group housed according to sex at a maximum of five mice per cage. We aimed to maintain an equal distribution of males and females; however, this was not always feasible due to the difficulty of reliably determining sex at the time of neonatal viral injection (P0-1). Group sizes and sex distributions for mice used for somatosensory and sensorimotor behaviour were as follows: neonatal (ctrl: n = 7 [3m/4f], hM3Dq: n = 8 [6m/2f]), juvenile (ctrl: n = 11 [5m/6f], hM3Dq: n = 14 [10m/4f]), and adolescent (ctrl: n = 7 [6m/1f], hM3Dq: n = 11 [4m/7f]). Neuropixels recordings and immunohistochemistry were performed in neonatal and juvenile cohorts, group sizes were: neonatal (ctrl: n = 9 [5m/3f], hM3Dq: n = 9 [4m/6f] and juvenile (ctrl: n = 7 [7m/1f], hM3Dq: n = 5 [3m/2f]). Animals lacking mCherry viral expression in dorsal root ganglia (DRG) and spinal cord were discarded from further analyses.

### Intraplantar viral injections

Unilateral intraplantar injections of a Cre-inducible adeno-associated viral vector encoding the excitatory hM3Dq DREADD (Designer Receptor Exclusively Activated by Designer Drugs; AAV9-hSyn-DIO-hM3D(Gq)-mCherry, #44361-AAV9, Addgene; working titre: 1.8×10^13^ vg/mL) or a control virus (AAV phSyn1(S)-FLEX-tdTomato-T2A-SypEGFP-WPRE, #51509-AAV1, Addgene; working titre: ≥ 4×10^12^ vg/mL) were administered at a volume of 2.5 μL into the left hind paw of newborn (P0-1). *TRPV1*^*Cre*^ transgenic mice (Wang et al. 2018). Injections were performed using a glass pipette attached to a 10 μL Hamilton syringe (7635-01, Hamilton Company). This design allowed us to compare the adult sensorimotor behavioural outcome of developmentally restricted transient primary afferent activation over one of our three putative critical periods.

Animals lacking mCherry viral expression in DRG and spinal cord were discarded from further analyses.

### CNO treatment and TRPV1+ primary afferent activation

Following the viral intraplantar injection, mice were randomly assigned to receive twice daily intraperitoneal (i.p.) injections of the DREADD ligand clozapine-N-oxide (CNO, 5 mg/kg; #4936, Tocris Bioscience) during one of three defined developmental windows: P8-12, P13-17, or P18-22. Both control and experimental animals received CNO injections. For each 5-day treatment period, CNO was administered twice daily with a 10-12 hr gap between injections (Kozorovitskiy et al. 2012; Salesse et al. 2020). After each injection, spontaneous behaviour (e.g. huddling; Supplementary Video V1) was monitored for 1 hr prior to mice being returned to their home cage. During the monitoring period, mice were kept in a heated holding box to maintain normal body temperature and were recorded using a GoPro HERO10 Black camera (60 frames per second (fps), 1080p resolution).

### Behavioural testing

Animals were acclimatised to the testing arena (7.5 × 7.5 × 15 cm transparent plastic chamber) for 60 minutes (min) per day over 2 to 3 days prior to experimentation and data collection. Between the ages of P30-37, mice underwent daily behavioural testing, following this sequence: 1) brush, 2) von Frey, 3) acetone cooling, 4) hot plate test, 5) pin prick, 6) catwalk, and 7) beam walking test.

#### Dynamic Touch Test (Brush)

To assess dynamic light touch sensitivity, mice were acclimatised as outlined above, and then habituated in the testing arena on an elevated wire grid (Ugo Basile, 0.6 mm grid width) for 30 min on the test day. The glabrous skin of the plantar hind paw surface was lightly stroked with a paintbrush from heel to toe. Each stroke was repeated five times, with 10 second (s) intervals between trials. The test included two rounds: the first round stimulated the hind paw ipsilateral to the injection (left hind paw), followed by a 5 min break before testing the contralateral side (right hind paw). Responses were scored as follows: 0 for no movement, 1 for brief paw lifting (~1 s or less), 2 for sustained lifting (more than 2 s), 3 for flinching or licking, and 4 for paw guarding (Duan et al. 2014). The average score from the five trials was used to quantify the touch response for each mouse. For developmental assessment of brush reflex sensitivity in pups, responses were scored as 0 - no response or 1 - withdrawal response, and sensitivity was expressed as the proportion of withdrawal responses over the total number of trials.

#### von Frey Test

To assess mechanical sensitivity to innocuous punctate stimuli, mice were placed on an elevated wire grid (Ugo Basile, 0.6 mm grid width) and habituated for 30 min on the test day. The plantar surface of the hind paw was stimulated with calibrated von Frey monofilaments (0.02-1 g). The test was conducted once per hind paw, starting with the ipsilateral side (left hind paw) with a 5 min interval between trials. The paw withdrawal threshold was determined on both left and right hind paws using Dixon’s up-down method (Dixon 1965).

#### Acetone cooling test

To assess cold sensitivity, mice were placed on an elevated wire grid (Ugo Basile, 0.6 mm grid width) and habituated for 30 min on the test day. A drop of acetone was gently applied to the plantar surface of the hind paw using a 5 mL syringe. The test not was conducted once per hind paw, with a 5 min interval between trials. Responses were scored as follows: 0 for no movement, 1 for brief paw lifting (~1 s or less), 2 for sustained lifting (more than 2 s), 3 for flinching or licking, and 4 for paw guarding.

#### Pin Prick test

Noxious punctate mechanical sensitivity was assessed using the Pin Prick test. On the test day, mice were placed on an elevated wire grid (Ugo Basile, 0.6 mm grid width) and habituated for 30 min. A 0.3 μL insulin syringe with a 30 gauge needle (Becton Dickinson GmbH) was then applied to the plantar surface of the hind paw without penetrating the skin. Responses were recorded over 3 trials, with a 1 min interval between trials. The test involved two rounds: the first round stimulated the hind paw ipsilateral to the injection (left hind paw), followed by a 5 min break before testing the contralateral side (right hind paw). Responses were scored as follows: 0 for no movement, 1 for brief paw lifting (~1 s or less), 2 for sustained lifting (more than 2 s), 3 for flinching or licking, and 4 for paw guarding. The average score from the three trials was used to measure the punctate mechanical sensitivity for each mouse.

#### Thermal sensitivity (Hot plate test)

Mice were individually placed on a hot plate (Ugo Basile) pre-heated to 30°C, with the temperature gradually increasing from 30°C to 50°C over the course of 5 min. A small heat-proof rescue platform was provided in the chamber, allowing the mice to escape the hot plate. Each mouse was video recorded using a GoPro HERO10 Black camera (60 fps, 1080 pixels resolution) for later offline analysis. The recordings were evaluated using BORIS software by an observer blinded to the experimental conditions. Thermally-evoked nociceptive behaviours, including hind paw licking/biting, toe tapping, and time spent on and off the platform, were quantified.

#### CatWalk

The CatWalk XT automated gait analysis system (Noldus Information Technology, Wageningen, The Netherlands) was used to assess gross locomotor function and motor coordination in adult mice. Mice were acclimated to the testing room for 30 min prior to the experiment. Three compliant runs (no more than 50% Run Maximum Variation) were recorded per mouse. During each run, the mouse was placed at one end of the CatWalk and allowed to walk across the walkway towards a darkened shelter. The following detection settings were used: camera gain = 19.58 dB, green intensity threshold = 0.1, red ceiling light = 17.9 V, and green walkway light = 16.9 V.

#### Beam walking test

Fine motor coordination was assessed using the beam walking assay. The beam consisted of 1-meter square shaped beam 1 cm in width, elevated 50 cm above the table on two poles. A black box at the end of the beam served as the end of the run, with a safety net placed underneath. Two markings on the beam, spaced 23 cm apart, indicated the trial distance. Mice were placed on a starting platform just outside the first marking and allowed to cross the beam at their own pace. Two infrared lamps illuminated the beam for video recording in the dark room. Video recordings were captured using a high-speed machine vision camera (Basler acA2000-340 kmNIR) at 280 fps, with an ISO of 3500 and a resolution of 250 × 1400 pixels. Each mouse was recorded crossing the 23 cm section of the beam twice in the right to left direction to quantify kinematics in the ipsilateral hindlimb. Compliant runs were defined as the mouse taking 3 consecutive steps within the 23 cm section. Two compliant runs were recorded and analysed. The start of a 3-step sequence was defined as the first frame in which the contralateral (right) hind paw initiated the swing phase while the ipsilateral (left) hind paw remained in stance phase; the sequence ended once the ipsilateral hind paw had completed three steps and the contralateral hind paw was poised to begin a new swing. Videos were later analysed using SLEAP (Pereira et al. 2022), a markerless pose estimation software as outlined below.

### SLEAP model training

A single-instance UNet-based network was trained using SLEAP with 2,085 manually labelled frames containing 14 anatomical landmarks (snout, neck, ipsilateral forepaw, contralateral forepaw, contralateral hind paw, ipsilateral hip, contralateral hip, ipsilateral knee, contralateral knee, ipsilateral calcaneum ankle, contralateral ankle, ipsilateral hind paw, contralateral hind paw, and tail base). Labelled frames were manually selected to represent the full range of step cycle phases (stance and swing). Training used a maximum stride of 16 and input scaling of 0.2, with standard data augmentation (random rotation, translation, flipping, and scaling) (Table S1). The validation set comprised 232 instances. For the final model the mean error distance was 5.19 pixels (1.04 mm), the mean Average Precision (mAP) was 0.70, and the mean Average Recall (mAR) was 0.74.

Per-frame 2D coordinates (x and y) for each body part were loaded from the CSV files. Missing samples were linearly interpolated and remaining edge gaps were filled (forward/backward fill). To reduce frame-to-frame jitter before kinematic calculations, each coordinate trace was smoothed with a Savitzky-Golay filter. For the ipsilateral hind paw, the Euclidean distance between consecutive frames was computed and summed to obtain total distance travelled over the analysis window. Instantaneous speed was computed per step as distance between frames divided by the known frame interval (Δt;1/280 s). Total time was the number of frames in the window multiplied by Δt, and overall speed was total distance divided by total time.

Angles were calculated from smoothed body part coordinates using vector geometry and the dot product. For the knee-ankle-paw (KAP) angle, vectors from the ankle to the hind paw (A to P) and from the ankle to the knee (A to K) were used; for the hip-knee-ankle (HKA) angle, vectors from the knee to the ankle (K to A) and from the knee to the hip (K to H) were used. Angles were computed as the arccosine of the normalised dot product and converted to degrees. Per-frame KAP and HKA angle values were extracted for each animal across two consecutive runs, with only animals present in both runs included for paired analysis. For each animal and run, angle distributions were generated as histograms (30 bins spanning 0-140° for KAP and 0-180° for HKA), and the total area under the curve (AUC) was calculated using the trapezoidal rule. To assess angle usage within specific ranges, histograms were subdivided into predefined intervals (KAP: 30-50°, 50-70°, 70-90°, 90-120°; HKA: 60-80°, 80-100°, 100-120°, 120-140°, 140-160°). The AUC for each interval was computed separately for the two runs and then averaged to produce per-animal, per-range values. These data were visualised as radar plots showing group means ± standard deviations.

### Neuropixels extracellular recordings

Mice were anaesthetised with an intraperitoneal injection of urethane (2 mg/g) and monitored every 10 min until areflexia was observed. Body temperature was maintained on a heating pad, and eye ointment was applied to prevent corneal damage. The dorsal area around the planned incision site was shaved, cleaned with 4% Chlorhexidine (Hibiscrub), and a longitudinal skin incision was made to expose the muscles and vertebral column at the lower thoracic to upper lumbar levels. To widen the surgical field, muscles and fascia along the spinal cord were cut. Without applying pressure to the spinal cord, a laminectomy was performed between thoracic (T) vertebra T13 and lumbar (L) vertebra L1 to expose the spinal cord. The thoracic vertebra immediately anterior to the laminectomy was secured with microscopic tweezers to stabilise the spinal cord during surgery and recordings.

Prior to insertion, the Neuropixels probe was coated with DiI (Cell Tracker CM-DiI, Invitrogen) diluted 1:1 in 100% ethanol. After gently removing the dura mater with a curved needle, the probe was lowered into the left hemisection of L4-5 to a depth of 500-700 μm to target the dorsal horn. Recordings were obtained using a Neuropixels 1.0 (3A) probe (Imec, Leuven, Belgium) (Jun et al. 2017) while the left (ipsilateral) hind paw was gently brushed with a soft paintbrush for 5-10 consecutive stimuli. At the end of the recording session, mice were transcardially perfused with formalin as described in Tissue Preparation and Immunohistochemistry, and spinal cord tissue was collected for histological analysis.

Spike sorting was performed with Kilosort4 (Pachitariu et al. 2024). Automated clusters were manually reviewed in Phy (https://github.com/cortex-lab/phy). Because all recordings were analysed at the multi-unit level, clusters were retained if they displayed consistent and stable spike waveforms clearly distinguishable from background noise. Clusters with <1.2 Hz firing rate were excluded from further analysis. No attempt was made to discriminate single units from multi-units; instead, all retained clusters were treated as multi-unit activity (MUA) for downstream analysis. To identify stimulus onsets, spike times and cluster assignments were loaded from Kilosort4 output and restricted to units within the dorsal horn depth range (0-500 μm) based on cluster depth information. Baseline activity for each unit was estimated during the 0-1 s pre-stimulus period (the 1 s preceding the five brush stimuli) by sliding a 50 ms counting window in 5 ms steps. The baseline mean and standard deviation (SD) of spike counts per unit defined the expected distribution. After 1 s, the same sliding window was advanced through the remainder of the recording, and units were marked as “active” if their firing exceeded a z-score of ≥3 relative to baseline, or if they produced ≥2 spikes when baseline variance was negligible. At the population level, the number of elevated units was tracked over time. A stimulus onset was defined when the proportion of active units exceeded 20% (and ≥3 units in absolute terms) continuously for 20-40 ms. To avoid spurious detections, a minimum spacing of 1.5 s was enforced between consecutive onsets, consistent with the expected inter-stimulus interval. This procedure identified five stimulus onsets per trial.

Raster plots and peri-stimulus time histograms (PSTHs) were then generated relative to these detected onsets. Spikes were aligned to each onset and displayed across trials, and PSTHs were computed in 20 ms bins as both raw firing rates (Hz) and z-scored responses. To prevent artificially inflated z-scores in low-rate units, a variance floor was applied: the baseline SD used for normalisation was constrained to be no smaller than either a Poisson-derived estimate of variance or a fixed minimum of 5 Hz. This ensured more stable and interpretable z-scores across the population. To classify stimulus-responsive clusters, clusters were labelled into three categories based on their mean z-score (mean-z) in the 0-1 s post-stimulus window: responders (mean-z ≥ 1), and non-responders (mean-z < 1).

### Anterograde tracing of S1 corticospinal input to the lumbar dorsal horn

Mice were placed in an induction chamber with 4% isoflurane in medical O_2_. Once induced, mice were positioned in a stereotaxic head frame (#51730U, Stoelting), and anaesthesia was maintained at 2% isoflurane in medical O_2_ via nosecone. Pups under the age of 15 days were induced in the stereotactic frame nosecone and placed in custom neonatal head bars. The head was shaved and preoperative analgesic EMLA cream (AstraZeneca) was applied on the skin overlaying the skull. Following disinfection with 4% Chlorhexidine (Hibiscrub), an incision was made in the skin overlaying the skull, and a craniotomy performed either via needle in pups, or via micro drill in adults (#58610, Stoelting). To trace the developing corticospinal projections from somatosensory cortex hindlimb region (S1hl), a Cre-dependent anterograde viral tracer (AAV9 hSyn-DIO-hM4D (Gi)-mCherry, #44362-AAV9, Addgene; working titre ≥ 1×10^13^ vg/mL) was injected in the S1hl of *Emx1*^*Cre*^ mice using a 1 μl Hamilton syringe (Model 7001 KH, Hamilton Company) attached to a manual microinjector (Model 5000, Kopf). The needle was left in place for 3 min after the injection to avoid leakage. Subsequently, the skin was sutured using Glueture (World Precision Instruments, US) and animals were monitored and allowed to recover in a heated incubator. Brains and spinal cords were processed for immunohistochemistry 7 days post injection, as outlined below. Sections were imaged with an inverted confocal microscope (SoRa Yokogawa CSU-W1 series, Nikon) and analysed using ImageJ software.

### Tissue Preparation and Immunohistochemistry

Spinal, peripheral and brain tissue were collected from all animals at the end of experiments to verify viral transfection. Mice in which Neuropixels recordings failed were still included in immunohistochemical analyses.

Mice were anaesthetised with pentobarbital (10 mg/g body weight) and transcardially perfused with 10% neutral buffered formalin (formalin solution, HT501640, Sigma-Aldrich). The spinal cord with attached DRGs was dissected and post-fixed overnight at 4 °C in the same fixative. Spinal cord tissue was then stored at 4 °C in 30% sucrose in phosphate buffer solution (PBS) containing 0.02% sodium azide (NaN_3_) until sectioning, while DRGs were embedded in optimal cutting temperature compound (O.C.T. Compound, Tissue-Tek) and stored at −70 °C to −80 °C until sectioning. Spinal cord sections (30 μm) were cut using a Leica microtome and stored at 4 °C in 5% sucrose with 0.02% NaN_3_. OCT-embedded DRGs were cryosectioned at 14 μm, mounted directly on Superfrost Plus adhesion microscope slides (J1800AMNZ, Espredia), and stored at −20 °C until staining. Prior to staining, sections were washed 3 times for 10 min each in cold 0.01 M PBS, permeabilised with PBT (PBS containing 0.1% Triton X-100), blocked for 1 hr at room temperature in PBT with 10% donkey serum, and incubated overnight at 4 °C with primary antibodies diluted in PBT containing 1% donkey serum. Primary antibody staining was detected using fluorophore-conjugated secondary antibodies. Images were acquired on a Zeiss LSM 800 confocal microscope. Quantitative analysis was performed in FIJI (Schindelin et al. 2012) on 2-5 sections per spinal cord, taken from segments L4 and L5.

The following primary antibodies were used: rabbit anti-DsRed (1:500, #632496, Takara), rabbit anti-PKCγ (1:1000, #sc-211, Santa Cruz), guinea pig anti-vGluT1 (1:1000, #ab5905, Millipore), sheep anti-CGRP (1:1000, #22560, Abcam), Alexa Fluor 488-conjugated isolectin B4 (IB4; 1:500, #I21411, Invitrogen), chicken anti-GFP (1:2000, #132006, Synaptic Systems) and guinea-pig anti-parvalbumin (1:1000, PV-GP-Af1000, Frontier Institute).

### Statistical Analysis

Statistical analyses were performed using Prism 10 (GraphPad Software). Scores recorded across repeated stimulations (brush, pinprick) and values from the CatWalk test were analysed using repeated-measures (RM) ANOVA. Post hoc comparisons were conducted using Sidak’s multiple comparisons test, and data are presented as mean ± SD. In the CatWalk test, “Limb” was treated as a within-subject factor, and no matching was applied. For between-group comparisons on individual sessions, average withdrawal scores were analysed using either two-tailed parametric (Student’s t-test) or non-parametric (Mann-Whitney test) statistics, as appropriate. These data are presented as box plots showing the median, with whiskers indicating minimum and maximum values. For the Neuropixels analysis, cluster-level response metrics (mean z-score values, depth) were averaged per animal to generate animal-level values for statistical testing. Group differences in mean-z were assessed using non-parametric Mann-Whitney tests. To further evaluate population responsiveness, clusters were classified as responders (mean-z ≥ 1) or non-responders (mean-z < 1), and their distributions compared between cohorts using Fisher’s exact tests. Cohort-level distributions by depth band were visualised with violin plots overlaid with individual animal points. The threshold for statistical significance was set at *P* < 0.05.

## Author Contributions

Conceptualization, L.A., S.C.K.; Data curation, L.A.; Formal analysis, L.A., C.J.B., S.C.K. Investigation, L.A., A.M.C., A.G., S.C.K.; Methodology, L.A., S.C.K. Project administration, L.A., S.C.K.; Resources, S.C.K.; Validation, L.A., C.J.B., S.C.K.; Visualization, L.A., S.C.K.; Writing – original draft, L.A., S.C.K.; Writing – reviewing and editing, L.A., S.C.K.

## Declarations

The authors declare no competing interests.

## Data availability

Data will be made available following reasonable request to S.C.K.

## Acknowledgements

We thank Dr. Daniel Bush (University College London) for discussions on Neuropixels analysis and Chloe Jensen for assistance with immunohistochemistry. This work was supported by Medical Research Foundation Fellowships MRF-087-0001-F-KOCH-C0917 (L.A.) and MRF-087-0003-F-KOCH-C0917 (S.C.K.), by MRF-160-0012-ELP-FABR-C0841 (A.C.), and by a Brain Research UK Miriam Marks Postdoctoral Research Fellowship in Neurodegenerative Diseases.

## References

Altman, J., and K. Sudarshan. 1975. ‘Postnatal development of locomotion in the laboratory rat’, Anim Behav, 23: 896–920.

Baccei, M. L., and M. Fitzgerald. 2004. ‘Development of GABAergic and glycinergic transmission in the neonatal rat dorsal horn’, J Neurosci, 24: 4749–57.

Barbeau, H., and S. Rossignol. 1987. ‘Recovery of locomotion after chronic spinalization in the adult cat’, Brain Res, 412: 84–95.

Beggs, S., G. Currie, M. W. Salter, M. Fitzgerald, and S. M. Walker. 2012. ‘Priming of adult pain responses by neonatal pain experience: maintenance by central neuroimmune activity’, Brain, 135: 404–17.

Beggs, S., C. Torsney, L. J. Drew, and M. Fitzgerald. 2002. ‘The postnatal reorganization of primary afferent input and dorsal horn cell receptive fields in the rat spinal cord is an activity-dependent process’, Eur J Neurosci, 16: 1249–58.

Belford, G. R., and H. P. Killackey. 1980. ‘The Sensitive Period in the Development of the Trigeminal System of the Neonatal Rat’, Journal of Comparative Neurology, 193: 335–50.

Boyle, K. A., M. A. Gradwell, T. Yasaka, A. C. Dickie, E. Polgar, R. P. Ganley, D. P. H. Orr, M. Watanabe, V. E. Abraira, E. D. Kuehn, A. L. Zimmerman, D. D. Ginty, R. J. Callister, B. A. Graham, and D. I. Hughes. 2019. ‘Defining a Spinal Microcircuit that Gates Myelinated Afferent Input: Implications for Tactile Allodynia’, Cell Rep, 28: 526–40 e6.

Carvell, G. E., and D. J. Simons. 1996. ‘Abnormal tactile experience early in life disrupts active touch’, J Neurosci, 16: 2750–7.

Cavanaugh, D. J., A. T. Chesler, J. M. Braz, N. M. Shah, D. Julius, and A. I. Basbaum. 2011. ‘Restriction of transient receptor potential vanilloid-1 to the peptidergic subset of primary afferent neurons follows its developmental downregulation in nonpeptidergic neurons’, J Neurosci, 31: 10119–27.

Cavanaugh, D. J., H. Lee, L. Lo, S. D. Shields, M. J. Zylka, A. I. Basbaum, and D. J. Anderson. 2009. ‘Distinct subsets of unmyelinated primary sensory fibers mediate behavioral responses to noxious thermal and mechanical stimuli’, Proc Natl Acad Sci U S A, 106: 9075–80.

Chang, E. F., and M. M. Merzenich. 2003. ‘Environmental noise retards auditory cortical development’, Science, 300: 498–502.

DiGiovanna, J., N. Dominici, L. Friedli, J. Rigosa, S. Duis, J. Kreider, J. Beauparlant, R. van den Brand, M. Schieppati, S. Micera, and G. Courtine. 2016. ‘Engagement of the Rat Hindlimb Motor Cortex across Natural Locomotor Behaviors’, J Neurosci, 36: 10440–55.

Dixon, W. J. 1965. ‘The up-and-down Method for Small Samples’, Journal of the American Statistical Association, 60: 967–78.

Donatelle, J. M. 1977. ‘Growth of the corticospinal tract and the development of placing reactions in the postnatal rat’, J Comp Neurol, 175: 207–31.

Duan, B., L. Cheng, S. Bourane, O. Britz, C. Padilla, L. Garcia-Campmany, M. Krashes, W. Knowlton, T. Velasquez, X. Ren, S. Ross, B. B. Lowell, Y. Wang, M. Goulding, and Q. Ma. 2014. ‘Identification of spinal circuits transmitting and gating mechanical pain’, Cell, 159: 1417–32.

Eccleston, C., and J. Clinch. 2007. ‘Adolescent chronic pain and disability: A review of the current evidence in assessment and treatment’, Paediatr Child Health, 12: 117–20.

Eccleston, C., S. Wastell, G. Crombez, and A. Jordan. 2008. ‘Adolescent social development and chronic pain’, Eur J Pain, 12: 765–74.

Feather-Schussler, D. N., and T. S. Ferguson. 2016. ‘A Battery of Motor Tests in a Neonatal Mouse Model of Cerebral Palsy’, J Vis Exp.

Fitzgerald, M. 2005. ‘The development of nociceptive circuits’, Nat Rev Neurosci, 6: 507-20.———. 2024. ‘On the relation of injury to pain-an infant perspective’, Pain, 165: S33–S38.

Fitzgerald, M., and S. Gibson. 1984. ‘The postnatal physiological and neurochemical development of peripheral sensory C fibres’, Neuroscience, 13: 933–44.

Fitzgerald, M., and E. Jennings. 1999. ‘The postnatal development of spinal sensory processing’, Proc Natl Acad Sci U S A, 96: 7719–22.

Fox, W. M. 1965. ‘Reflex-ontogeny and behavioural development of the mouse’, Anim Behav, 13: 234–41.

Franks, K. M., and J. S. Isaacson. 2005. ‘Synapse-specific downregulation of NMDA receptors by early experience: a critical period for plasticity of sensory input to olfactory cortex’, Neuron, 47: 101–14.

Gorski, J. A., T. Talley, M. Qiu, L. Puelles, J. L. Rubenstein, and K. R. Jones. 2002. ‘Cortical excitatory neurons and glia, but not GABAergic neurons, are produced in the Emx1-expressing lineage’, J Neurosci, 22: 6309–14.

Goulding, M., S. Bourane, L. Garcia-Campmany, A. Dalet, and S. Koch. 2014. ‘Inhibition downunder: an update from the spinal cord’, Curr Opin Neurobiol, 26: 161–6.

Granmo, M., P. Petersson, and J. Schouenborg. 2008. ‘Action-based body maps in the spinal cord emerge from a transitory floating organization’, J Neurosci, 28: 5494–503.

Grillner, S. 2006. ‘Biological pattern generation: the cellular and computational logic of networks in motion’, Neuron, 52: 751–66.

Henry, R. J., V. E. Meadows, B. A. Stoica, A. I. Faden, and D. J. Loane. 2020. ‘Longitudinal Assessment of Sensorimotor Function after Controlled Cortical Impact in Mice: Comparison of Beamwalk, Rotarod, and Automated Gait Analysis Tests’, J Neurotrauma, 37: 2709–17.

Hensch, T. K. 2004. ‘Critical period regulation’, Annual Review of Neuroscience, 27: 549–79.

Holmberg, H., and J. Schouenborg. 1996. ‘Postnatal development of the nociceptive withdrawal reflexes in the rat: a behavioural and electromyographic study’, J Physiol, 493 (Pt 1): 239–52.

Hsu, J. Y., S. A. Stein, and X. M. Xu. 2006. ‘Development of the corticospinal tract in the mouse spinal cord: a quantitative ultrastructural analysis’, Brain Res, 1084: 16–27.

Hubel, D. H., and T. N. Wiesel. 1970. ‘The period of susceptibility to the physiological effects of unilateral eye closure in kittens’, J Physiol, 206: 419–36.

Jun, J. J., N. A. Steinmetz, J. H. Siegle, D. J. Denman, M. Bauza, B. Barbarits, A. K. Lee, C. A. Anastassiou, A. Andrei, C. Aydin, M. Barbic, T. J. Blanche, V. Bonin, J. Couto, B. Dutta, S. L. Gratiy, D. A. Gutnisky, M. Hausser, B. Karsh, P. Ledochowitsch, C. M. Lopez, C. Mitelut, S. Musa, M. Okun, M. Pachitariu, J. Putzeys, P. D. Rich, C. Rossant, W. L. Sun, K. Svoboda, M. Carandini, K. D. Harris, C. Koch, J. O’Keefe, and T. D. Harris. 2017. ‘Fully integrated silicon probes for high-density recording of neural activity’, Nature, 551: 232–36.

Keller, A. F., J. A. Coull, N. Chery, P. Poisbeau, and Y. De Koninck. 2001. ‘Region-specific developmental specialization of GABA-glycine cosynapses in laminas I-II of the rat spinal dorsal horn’, J Neurosci, 21: 7871–80.

Knudsen, E. I. 1985. ‘Experience Alters the Spatial Tuning of Auditory Units in the Optic Tectum during a Sensitive Period in the Barn Owl’, Journal of Neuroscience, 5: 3094–109.

Koch, S. C. 2019. ‘Motor task-selective spinal sensorimotor interneurons in mammalian circuits’, Current Opinion in Physiology, 8.

Koch, S. C., M. G. Del Barrio, A. Dalet, G. Gatto, T. Gunther, J. Zhang, B. Seidler, D. Saur, R. Schule, and M. Goulding. 2017. ‘RORbeta Spinal Interneurons Gate Sensory Transmission during Locomotion to Secure a Fluid Walking Gait’, Neuron, 96: 1419–31 e5.

Koch, S. C., and M. Fitzgerald. 2013. ‘Activity-dependent development of tactile and nociceptive spinal cord circuits’, Ann N Y Acad Sci, 1279: 97–102.

Koch, S. C., K. K. Tochiki, S. Hirschberg, and M. Fitzgerald. 2012. ‘C-fiber activity-dependent maturation of glycinergic inhibition in the spinal dorsal horn of the postnatal rat’, Proc Natl Acad Sci U S A, 109: 12201–6.

Kozorovitskiy, Y., A. Saunders, C. A. Johnson, B. B. Lowell, and B. L. Sabatini. 2012. ‘Recurrent network activity drives striatal synaptogenesis’, Nature, 485: 646–50.

Liu, Y., A. Latremoliere, X. Li, Z. Zhang, M. Chen, X. Wang, C. Fang, J. Zhu, C. Alexandre, Z. Gao, B. Chen, X. Ding, J. Y. Zhou, Y. Zhang, C. Chen, K. H. Wang, C. J. Woolf, and Z. He. 2018. ‘Touch and tactile neuropathic pain sensitivity are set by corticospinal projections’, Nature, 561: 547–50.

Martin, J. H. 2005. ‘The corticospinal system: from development to motor control’, Neuroscientist, 11: 161–73.

Martin, J. H., M. Choy, S. Pullman, and Z. Meng. 2004. ‘Corticospinal system development depends on motor experience’, J Neurosci, 24: 2122–32.

Micheva, K. D., and C. Beaulieu. 1995. ‘Neonatal sensory deprivation induces selective changes in the quantitative distribution of GABA-immunoreactive neurons in the rat barrel field cortex’, J Comp Neurol, 361: 574–84.

Miraucourt, L. S., R. Dallel, and D. L. Voisin. 2007. ‘Glycine inhibitory dysfunction turns touch into pain through PKCgamma interneurons’, PLoS One, 2: e1116.

Neumann, S., J. M. Braz, K. Skinner, I. J. Llewellyn-Smith, and A. I. Basbaum. 2008. ‘Innocuous, not noxious, input activates PKCgamma interneurons of the spinal dorsal horn via myelinated afferent fibers’, J Neurosci, 28: 7936–44.

Pachitariu, M., S. Sridhar, J. Pennington, and C. Stringer. 2024. ‘Spike sorting with Kilosort4’, Nat Methods, 21: 914–21.

Pereira, T. D., N. Tabris, A. Matsliah, D. M. Turner, J. Li, S. Ravindranath, E. S. Papadoyannis, E. Normand, D. S. Deutsch, Z. Y. Wang, G. C. McKenzie-Smith, C. C. Mitelut, M. D. Castro, J. D’Uva, M. Kislin, D. H. Sanes, S. D. Kocher, S. S. Wang, A. L. Falkner, J. W. Shaevitz, and M. Murthy. 2022. ‘SLEAP: A deep learning system for multi-animal pose tracking’, Nat Methods, 19: 486–95.

Petersson, P., M. Granmo, and J. Schouenborg. 2004. ‘Properties of an adult spinal sensorimotor circuit shaped through early postnatal experience’, Journal of Neurophysiology, 92: 280–8.

Petitjean, H., S. A. Pawlowski, S. L. Fraine, B. Sharif, D. Hamad, T. Fatima, J. Berg, C. M. Brown, L. Y. Jan, A. Ribeiro-da-Silva, J. M. Braz, A. I. Basbaum, and R. Sharif-Naeini. 2015. ‘Dorsal Horn Parvalbumin Neurons Are Gate-Keepers of Touch-Evoked Pain after Nerve Injury’, Cell Rep, 13: 1246–57.

Salesse, C., J. Charest, H. Doucet-Beaupre, A. M. Castonguay, S. Labrecque, P. De Koninck, and M. Levesque. 2020. ‘Opposite Control of Excitatory and Inhibitory Synapse Formation by Slitrk2 and Slitrk5 on Dopamine Neurons Modulates Hyperactivity Behavior’, Cell Rep, 30: 2374–86 e5.

Schindelin, J., I. Arganda-Carreras, E. Frise, V. Kaynig, M. Longair, T. Pietzsch, S. Preibisch, C. Rueden, S. Saalfeld, B. Schmid, J. Y. Tinevez, D. J. White, V. Hartenstein, K. Eliceiri, P. Tomancak, and A. Cardona. 2012. ‘Fiji: an open-source platform for biological-image analysis’, Nat Methods, 9: 676–82.

Terashima, T. 1995. ‘Anatomy, development and lesion-induced plasticity of rodent corticospinal tract’, Neurosci Res, 22: 139–61.

Ueno, M., Y. Nakamura, J. Li, Z. Gu, J. Niehaus, M. Maezawa, S. A. Crone, M. Goulding, M. L. Baccei, and Y. Yoshida. 2018. ‘Corticospinal Circuits from the Sensory and Motor Cortices Differentially Regulate Skilled Movements through Distinct Spinal Interneurons’, Cell Rep, 23: 1286–300 e7.

Waldenstrom, A., M. Christensson, and J. Schouenborg. 2009. ‘Spontaneous movements: Effect of denervation and relation to the adaptation of nociceptive withdrawal reflexes in the rat’, Physiol Behav, 98: 532–6.

Walker, S. M. 2014. ‘Neonatal pain’, Paediatr Anaesth, 24: 39–48.

Walker, S. M., L. S. Franck, M. Fitzgerald, J. Myles, J. Stocks, and N. Marlow. 2009. ‘Long-term impact of neonatal intensive care and surgery on somatosensory perception in children born extremely preterm’, Pain, 141: 79–87.

Wang, F., E. Belanger, S. L. Cote, P. Desrosiers, S. A. Prescott, D. C. Cote, and Y. De Koninck. 2018. ‘Sensory Afferents Use Different Coding Strategies for Heat and Cold’, Cell Rep, 23: 2001–13.

Wiesel, T. N., and D. H. Hubel. 1963. ‘Single-Cell Responses in Striate Cortex of Kittens Deprived of Vision in 1 Eye’, Journal of Neurophysiology, 26: 1003–+.

Xu, Y., S. C. Koch, A. Chamessian, Q. He, M. Sundukova, P. Heppenstall, R. Ji, M. Fitzgerald, and S. Beggs. 2024. ‘Microglial Refinement of A-Fiber Projections in the Postnatal Spinal Cord Dorsal Horn Is Required for Normal Maturation of Dynamic Touch’, J Neurosci, 44.

Yoo, J. J., E. K. Serafin, J. M. Kofron, and M. L. Baccei. 2025. ‘Early life injury alters spinal astrocyte development’, J Neurosci.

