## Supplementary Material for "Discrete and sequential critical periods organise the development of task-specific sensorimotor circuits in mice"

**Supplementals**

*Histological characterization of CGRP⁺ and IB4⁺ afferent terminals in the dorsal horn*

For the CGRP/IB4 analysis, four to five transverse spinal cord sections corresponding to segments L4/L5 were selected per animal. $CGRP^+$ and $IB4^+$ areas were quantified in each hemisection individually using FIJI (Schindelin et al. 2012). Images were cropped to include only the dorsal horn, with the lower boundary defined by the central canal. After converting images to 8-bit, channels were split to allow separate analysis of CGRP and IB4. A default threshold was applied to generate selection areas, from which the positive area (in pixels) was measured. Measurements from left and right hemisections were averaged so that each animal was represented by a single value derived from four to five sections. Data are shown as box plots displaying the median and full range. Immunohistochemistry were performed in neonatal and juvenile cohorts, group sizes were: neonatal (ctrl: n = 4, hM3Dq: n = 7), juvenile (ctrl: n = 4, hM3Dq: n = 9). Group differences in $CGRP^+$ and $IB4^+$ area were assessed using non-parametric Mann-Whitney tests, with significance set at $P < 0.05$.

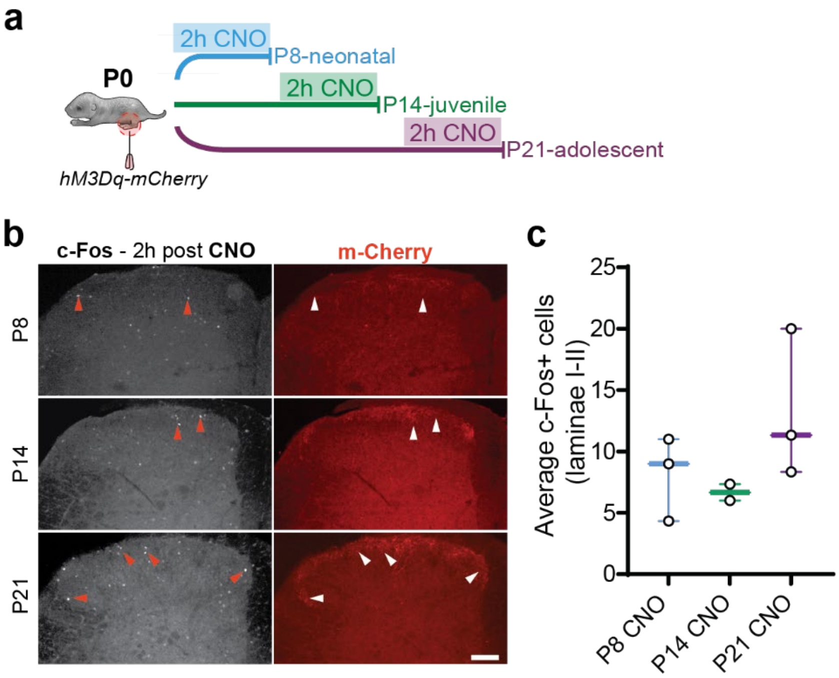


**Fig. S1.** Functional expression of hM3Dq confirmed by c-Fos activation in primary afferent termination zones following systemic CNO administration.

(a) Schematic of the experimental design. AA9-hM3Dq-mCherry was injected intraplantarly at P0, and acute CNO was administered i.p. at P8, P14, or P21. (b) Representative images showing hM3Dq-mCherry expression in the ipsilateral L4/L5 dorsal horn at (top to bottom) 8, 14, and 21 days post-injection, together with c-Fos expression 2 hr after CNO administration (scale bar 100 µm). (c) Quantification of c-Fos⁺ cells in the ipsilateral L4/L5 dorsal horn at each developmental stage (Kruskal-Wallis test, *n.s.*, n = 2-3 per age).

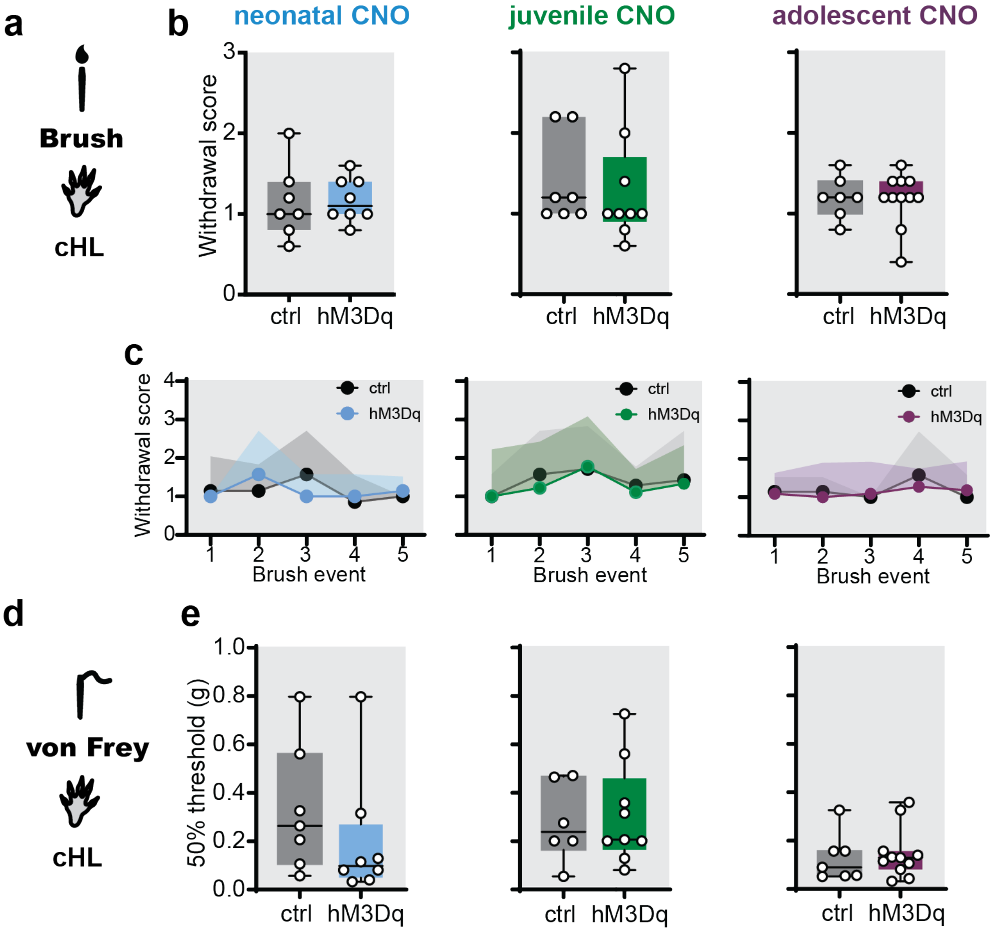


**Fig. S2.** No long-term effects of early-life manipulation on adult contralateral tactile sensitivity.

(a) Schematic of brush stimulation of the contralateral hind paw. (b) Average withdrawal to five consecutive dynamic brush strokes of the contralateral hind paw was unchanged in neonatal (left; P8-12), juvenile (middle; P13-17), and adolescent (right; P18-22) groups (unpaired t-tests or Mann-Whitney tests, as appropriate). (c) Trial-by-trial responses to individual brush strokes were likewise unchanged across groups (all *n.s.*, two-way ANOVA). (d) Schematic of von Frey hair stimulation of the contralateral hind paw. (e) Static touch sensitivity (50% von Frey threshold) was also unaffected in neonatal (left), juvenile (middle), and adolescent (right) mice. Group sizes: neonatal, n = 7 ctrl, 8 hM3Dq; juvenile, n = 7 ctrl, 9 hM3Dq; adolescent, n = 7 ctrl, 11 hM3Dq. TRPV1Cre mice expressing hM3Dq are shown in blue (neonatal), green (juvenile), and purple (adolescent); controls are shown in grey. All testing was performed in adulthood following transient developmental primary afferent activation.

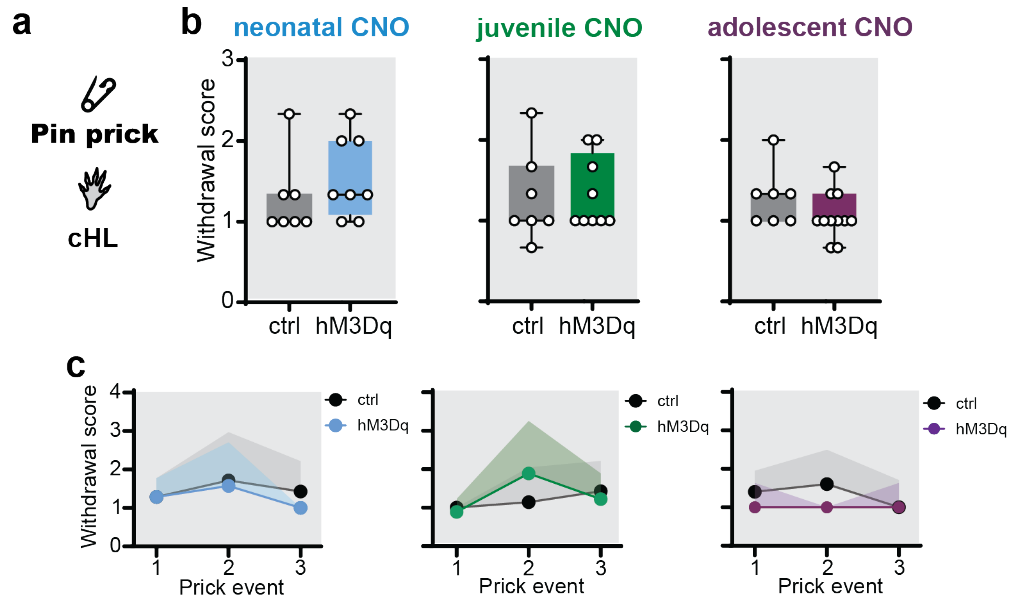


**Fig. S3.** Adult contralateral hind paw withdrawal responses to pin prick are unchanged by early-life manipulation.

(a) Schematic of pin prick stimulation of the contralateral hind paw. (b) Average withdrawal scores to three pin-prick stimuli in the contralateral hind paw was unchanged in neonatal (left; P8-12), juvenile (middle; P13-17), and adolescent (right; P18-22) groups (unpaired t-tests or Mann–Whitney tests, as appropriate). (c) Trial-by-trial withdrawal responses were likewise unchanged. RM two-way ANOVA confirmed no significant group or stimulus effects in neonatal (left), juvenile (middle), or adolescent (right) groups. Group sizes: neonatal, n = 7 ctrl, 8 hM3Dq; juvenile, n = 7 ctrl, 9 hM3Dq; adolescent, n = 7 ctrl, 11 hM3Dq. *TRPV1*$^{Cre}$ mice expressing hM3Dq are shown in blue (neonatal), green (juvenile), and purple (adolescent); controls in grey. All testing was performed in adulthood following transient developmental primary afferent activation.

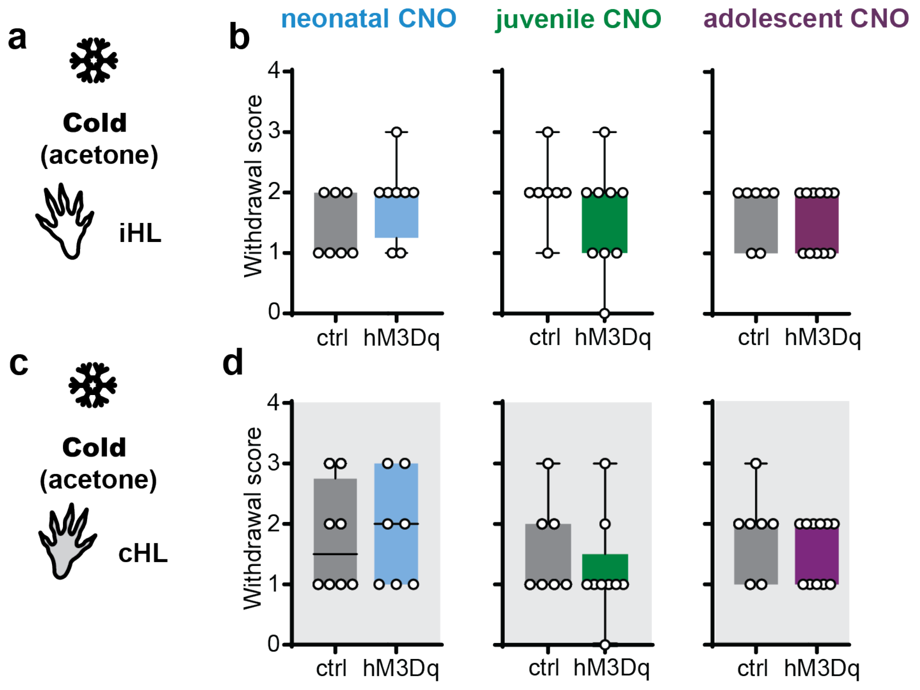


**Fig. S4.** Adult hind paw responses to noxious cold are unaffected by early-life afferents manipulation.

(a) Schematic of cold stimulation of the ipsilateral hind paw. (b) Withdrawal scores in response to acetone application on the plantar surface of the ipsilateral hind paw remained unchanged in neonatal (left; P8-12), juvenile (middle; P13-17) and adolescent (right; P18-22) groups (unpaired t-test or Mann-Whitney test, as appropriate). (a) Schematic of cold stimulation of the contralateral hind paw. (d) No significant differences were observed in the withdrawal amplitude scores from the contralateral hind paw (all *n.s.*, unpaired t-test or Mann-Whitney test, as appropriate). Group sizes: neonatal, n = 7 ctrl, 8 hM3Dq; juvenile, n = 6/7 ctrl, 9 hM3Dq; adolescent, n = 7 ctrl, 11 hM3Dq. *TRPV1*$^{Cre}$ mice expressing hM3Dq are shown in blue (neonatal), green (juvenile), and purple (adolescent); controls are shown in grey. Panels in the shaded grey background show contralateral hind paw responses. All testing was performed in adulthood following transient developmental primary afferent activation.

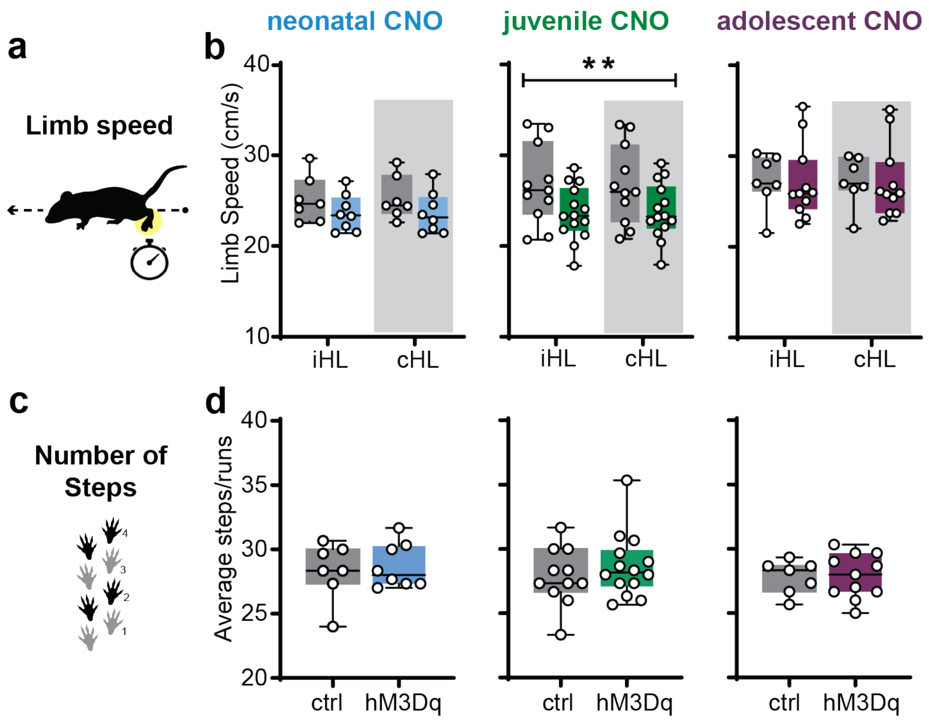


**Fig. S5**. Early-life manipulation reduces adult limb speed in juveniles but not in neonates or adolescents.

(a) Schematic of limb speed measurement on the CatWalk. (b) Limb speed was unaffected in the neonatal (left; P8-12) and in the adolescent (right; P18-22) group. In contrast, juveniles (middle; P18-22) displayed a reduction in limb speed (B; Group: $F_{1,46}$ = 8.278, $P$ = 0.01; Limb: $F_{1,46}$ = 0.01297, $P$ > 0.05; Interaction: $F_{1,46}$ = 0.01024, $P$ > 0.05). (c) Schematic of step count on the CatWalk. (d) The average number of steps was not significantly different between groups at any age (all *n.s.*, unpaired t-tests). Group sizes: neonatal, n = 7 ctrl, 8 hM3Dq; juvenile, n = 11 ctrl, 14 hM3Dq; adolescent, n = 7 ctrl, 11 hM3Dq. *TRPV1*$^{Cre}$ mice expressing hM3Dq are shown in blue (neonatal), green (juvenile), and purple (adolescent); controls in grey. Contralateral hindlimb responses are displayed on a shaded grey background. All testing was performed in adulthood following transient developmental primary afferent activation.

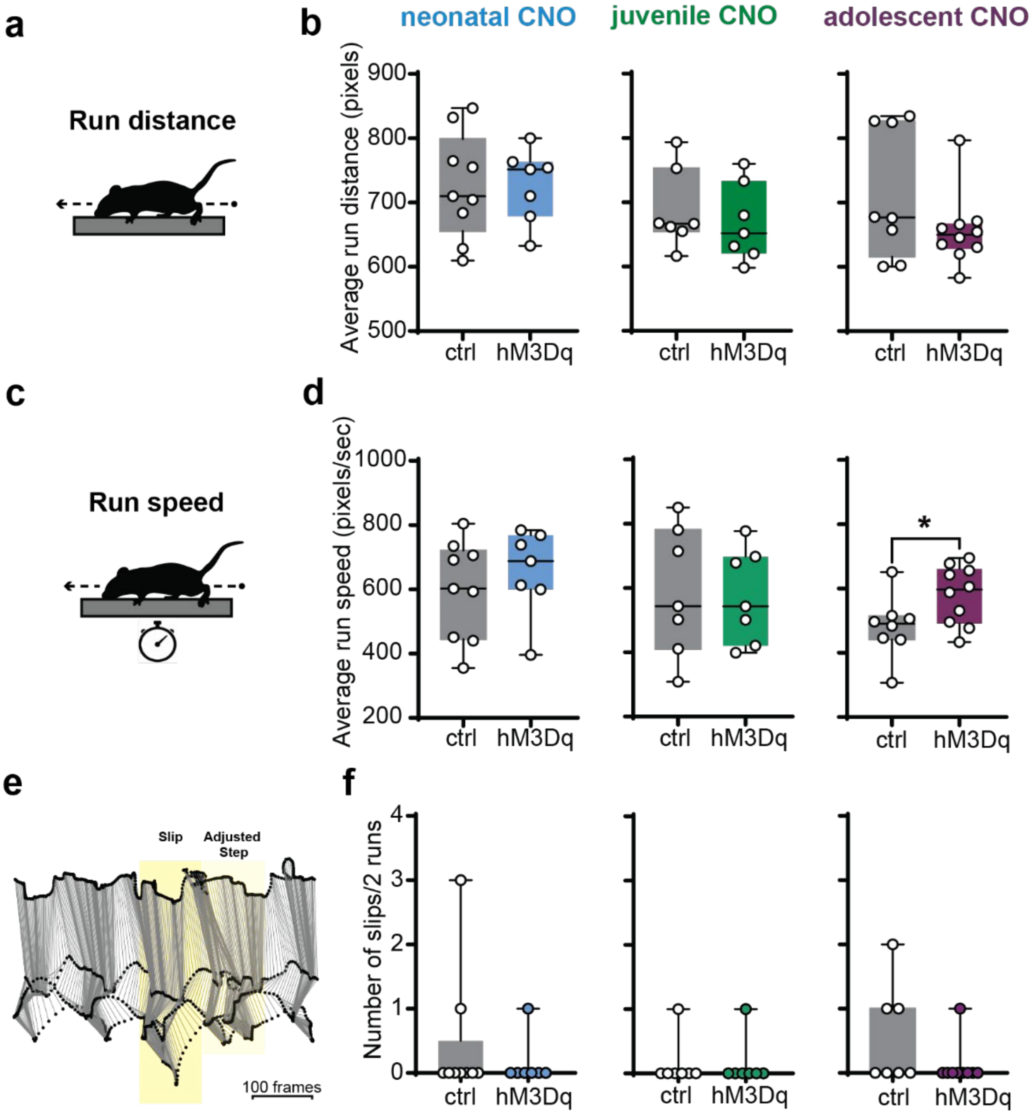


**Fig. S6.** Adult run speed is reduced in adolescents, but distance travelled, and slips are unaffected by early-life manipulation.

(a) Schematic of distance travelled on the beam. (b) Similar distance travelled were measured in neonatal (left; P8-12), juvenile (middle; P13-17) and adolescent (right; P18-22) (all *n.s.*, unpaired t-test) mice. (c) Schematic of run speed measurement on the beam. (d) Run speed was unchanged in the neonatal (left) and juvenile (middle; both *n.s.*, unpaired t-tests) mice but reduced in the adolescent group (right; ctrl vs hM3Dq, $^*P < 0.05$, unpaired t-tests). (e) Representative stick figure diagrams showing three complete hindlimb step cycles (swing and stance). The light-yellow shaded area in the representative limb traces during three steps indicates two step adjustments, while the dark yellow shaded area highlights a hind paw slip. (f) The number of slips across two three-step runs did not differ between groups (all *n.s.*, unpaired t-test) . Group sizes: neonatal, n = 9 ctrl, 7 hM3Dq; juvenile, n = 7 ctrl, 7 hM3Dq; adolescent, n = 7 ctrl, 10 hM3Dq. *TRPV1*$^{Cre}$ mice expressing hM3Dq are shown in blue (neonatal), green (juvenile), and purple (adolescent), controls in grey. All testing was performed in adulthood following transient developmental primary afferent activation.

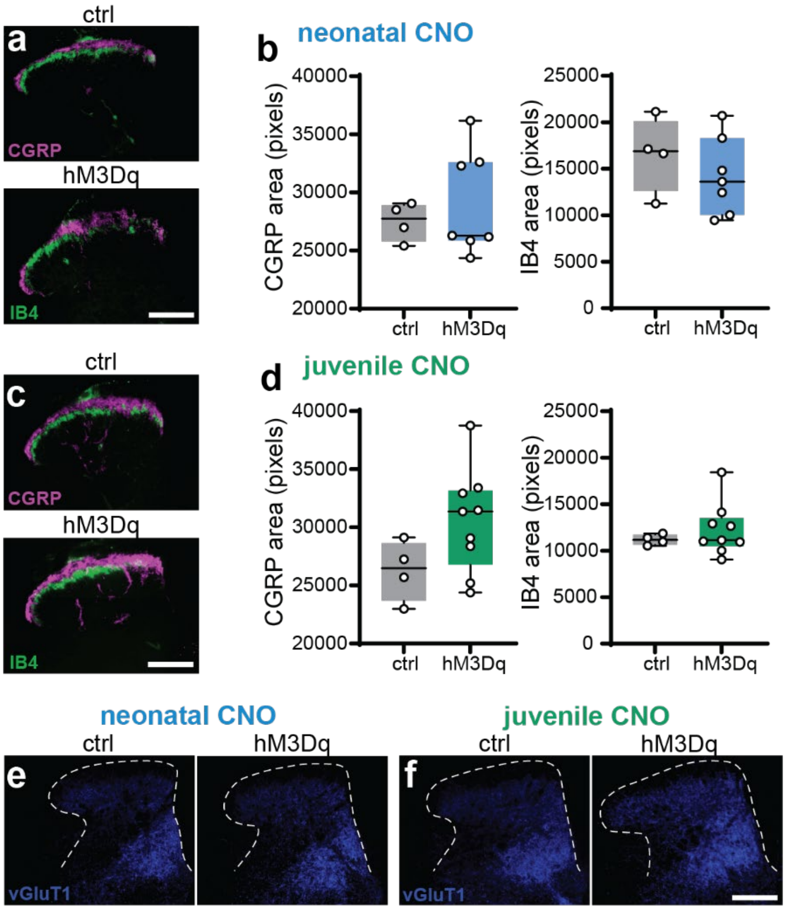


**Fig. S7.** CGRP⁺, IB4⁺, or vGluT1⁺ terminal distribution in the dorsal horn following early-life TRPV1+ afferent activation.

(a) Representative spinal cord sections immunostained for CGRP⁺ (magenta) and IB4⁺ (green) afferent terminals in the ipsilateral dorsal horn of neonatally (P8-12) treated mice. (b) Quantification of CGRP⁺ and IB4⁺ terminal area in ipsilateral and contralateral dorsal horn showed no significant differences between groups (all *n.s.*, unpaired t-tests). (c) Representative spinal cord sections immunostained for CGRP⁺ (magenta) and IB4⁺ (green) afferent terminals in the ipsilateral dorsal horn of juvenile (P13-17) treated mice. (d) Quantification of CGRP⁺ and IB4⁺ terminal area in ipsilateral and contralateral dorsal horn showed no significant differences between groups (all *n.s.*, unpaired t-tests). (e) Representative images of vGluT1⁺ afferent terminals in the ipsilateral dorsal horn of neonatally treated mice. (f) Representative images of vGluT1⁺ afferent terminals in the ipsilateral dorsal horn of juvenile treated mice. Scale bar: 200 µm.

**Video V1.** Spontaneous behaviour of CNO-treated mice at different developmental stages. Twenty-second clips show P8, P13, and P18 mice recorded 15 min after systemic CNO administration. Activation of TRPV1⁺ afferents during development did not alter spontaneous behaviours such as huddling, where pups remain in close physical contact and cluster together; a typical behaviour reflecting thermoregulation and social bonding.

**Table 1.** Training parameters for the SLEAP model. Key hyperparameters and augmentation settings used for single-instance SLEAP training, including network architecture, optimization, and data augmentation parameters.

| Parameter | Value |
|---|---|
| Max stride | 16 |
| Filters | 16 |

| | |
|---|---|
| Filters rate | 2 |
| Sigma | 2 |
| Output stride | 2 |
| **Data** | |
| Input scaling | 0.2 |
| **Optimization** | |
| Batch size | 4 |
| Epoch | 200 |
| Initial learning rate | 0.0001 |
| **Augmentation** | |
| Rotation minimum angle | -15 |
| Rotation maximum angle | 15 |
| Gaussian noise mean | 5 |
| Gaussian noise standard deviation | 1 |
| Contrast minimum gamma | 0.5 |
| Contrast maximum gamma | 2 |
| Brightness minimum value | 0 |
| Brightness maximum value | 10 |

### References


Schindelin, J., I. Arganda-Carreras, E. Frise, V. Kaynig, M. Longair, T. Pietzsch, S. Preibisch, C. Rueden, S. Saalfeld, B. Schmid, J. Y. Tinevez, D. J. White, V. Hartenstein, K. Eliceiri, P. Tomancak, and A. Cardona. 2012. 'Fiji: an open-source platform for biological-image analysis', *Nat Methods*, 9: 676-82.
